# Atypical B cells mediate poor response to Bacillus Calmette Guérin immunotherapy in non-muscle invasive bladder cancer

**DOI:** 10.1101/2022.12.30.522127

**Authors:** Priyanka Yolmo, Sadaf Rahimi, Stephen Chenard, Gwenaëlle Conseil, Danielle Jenkins, Kartik Sachdeva, Isaac Emon, Jake Hamilton, Minqi Xu, Manu Rangachari, Eva Michaud, Jose Mansure, Wassim Kassouf, David M. Berman, D. Robert Siemens, Madhuri Koti

**Author notes:** Equal Contribution (Priyanka Yolmo, Sadaf Rahimi).

## Abstract

Poor response to Bacillus Calmette-Guérin (BCG) immunotherapy remains a major barrier in the management of patients with non-muscle-invasive bladder cancer (NMIBC). Among the multiple factors contributing to poor outcomes, a B cell infiltrated pre-treatment immune microenvironment of NMIBC tumors has emerged as a key determinant of response to BCG. The mechanisms underlying the paradoxical roles of B cells in NMIBC are poorly understood. Here, we show that B cell dominant tertiary lymphoid structures (TLSs), a hallmark feature of chronic mucosal immune response, are abundant and located close to the epithelial compartment in pre-treatment tumors from BCG non-responders. Digital spatial proteomic profiling of whole tumor sections revealed higher expression of immune exhaustion-associated proteins within the TLSs from both responders and non-responders. Chronic local inflammation, induced by the N-butyl- N-(4-hydroxybutyl) nitrosamine (BBN) carcinogen, led to TLS formation with recruitment and differentiation of the immunosuppressive atypical B cell (ABCs) subset within the bladder microenvironment, predominantly in aging female mice compared to their male counterparts. Depletion of ABCs simultaneous to BCG treatment delayed cancer progression in female mice. Our findings provide the first evidence indicating the role of ABCs in BCG response and will inform future development of therapies targeting the B cell exhaustion axis.

## Introduction

Bladder cancer is the most common urological malignancy known worldwide with 573,278 cases diagnosed in 2020^1^. The majority of bladder cancer patients (75-85%) are diagnosed with early-stage NMIBC with a median age at diagnosis of 73 years. Exposure to environmental and occupational carcinogens, tobacco smoking, recurrent urinary tract infections, and aging can be attributed as the major risk factors for bladder cancer.^2^ Although more common in males,^3^ females with bladder cancer generally present with advanced-stage tumors and experience shorter recurrence-, and progression-free survival^4–7^. Following transurethral resection of bladder tumor (TURBT), patients categorized as intermediate or high-risk NMIBC are treated with intravesical Bacillus Calmette-Guérin (BCG) immunotherapy^8^. The treatment regimen involves an induction phase of 6 weekly instillations after tumor resection, followed by a maintenance phase of 3 weekly instillations every 3-6 months for over 1-3 years depending on the risk stratification^9^. BCG treatment in NMIBC patients is linked to a decreased risk of recurrence and progression when compared to transurethral resection alone. Despite its proven efficacy, recurrence post-BCG treatment occurs in over 50% of patients, necessitating repeated treatments, continuous surveillance, and cystectomy upon progression to muscle-invasive stage^10,11^.

Our previous study discovered a significant association between increased density of intra-tumoral B cells and poor outcomes in patients with NMIBC^12^. Tumor-infiltrating B cells are mainly localized within stromal aggregates of immune cells called tertiary lymphoid structures (TLSs)^13^. TLS formation in the bladder mucosa is induced due to chronic inflammation,^14^ such as persistent urinary tract infection^15,16^; exposure to carcinogens or smoking; immunomodulatory therapy^13^; or age-related increase in systemic levels of tumor necrosis factor-α (TNF-α)^17,18^. Indeed, our previous report established an increase in TLS density and maturation with advanced stages of bladder cancer^19^.

Simultaneous to biological aging-associated decline in the periphery, B cell proportions significantly increase at mucosal sites, predominantly within mucosa-associated TLSs^17,20–24^. Biological aging, repeated vaccination, and chronic inflammation also lead to the expansion of a circulating B cell subset known as atypical B cells (ABCs). ABCs are observed at a significantly higher frequency in females compared to males, exhibit features of high self-reactivity, and are equipped with specialized antigen-presenting ability^24–28^. In addition, ABCs are known to produce high levels of IL-6 and TNF-α and augment the suppression of B cell lymphopoiesis^29–31^. In aging mice, ABCs increase in the bone marrow with a more pronounced expansion in the spleen, produce autoantibodies and respond uniquely to innate/microbial stimuli^26,32^. A key feature of ABCs is their proliferative response to stimuli that activate endosomal nucleic acid sensing pathways (toll-like receptors; TLR7/9) leading to the production of immunosuppressive cytokines such as IL-10^26,33–36^.

Following intravesical instillation, BCG antigens are internalized by residual cancer cells, normal urothelial cells, and immune cells, leading to immune activation *via* various cytosolic and cell surface receptors such as TLR2/9^36–38^. We hypothesized that repeated local exposure to BCG during the induction phase of the treatment, potentiates systemic expansion of ABCs and recruitment to the bladder. Local immunosuppression mediated by ABCs potentially contributes to disease progression. In this study, we first characterized TLSs in pre-treatment tumors from patients, categorized as BCG responders and non-responders, using multiplex immunofluorescence (mIF) and NanoString GeoMx digital spatial technology (DSP). Since B cells/ABCs exhibit a sex and age-dependent expansion and response to persistent immune activation, we further investigated the potential role of ABCs in BBN carcinogen (mimicking carcinogen present in tobacco smoking) exposed aging wild type C57BL/6 and the four-core genotype (FCG) mice treated with BCG. Overall, our study is the first to demonstrate the expansion of ABCs following chronic carcinogen exposure and repeated BCG immunotherapy as one of the underlying factors for poor response to BCG immunotherapy, specifically, in females.

## RESULTS

### Pre-treatment tumor adjacent TLSs are increased in BCG non-responders and express proteins reflective of immune exhaustion

Based on our previous observation of high stromal B cell density and its association with early recurrence and progression, we questioned whether pre-treatment TLSs (characterized by CD79a+ B cells, CD3+ and CD8+ T cells, CD208+ mature dendritic cells (DCs), CD21+ follicular DCs and PNAd+ high endothelial venules) associate with response to BCG. Histopathological evaluation of BCG-naïve whole tumor sections from 28 patients (clinical details in **Suppl. Table S1**), revealed the presence of both small and large immune cell aggregates within the lamina propria, indicative of chronic inflammation in the bladder mucosa (**Fig. 1A**). To determine the maturation stage of TLSs, we performed mIF staining, which revealed varying organization and maturation stages of TLSs within the tumors from both BCG responders and non-responders (**Fig. 1B**). Overall, lymphoid aggregates and mature TLSs (with PNAd+ high endothelial venules, CD21+CD208+ DCs) were higher in whole tumor sections from BCG non-responders compared to responders (**Fig. 1C**).

**Figure 1.**
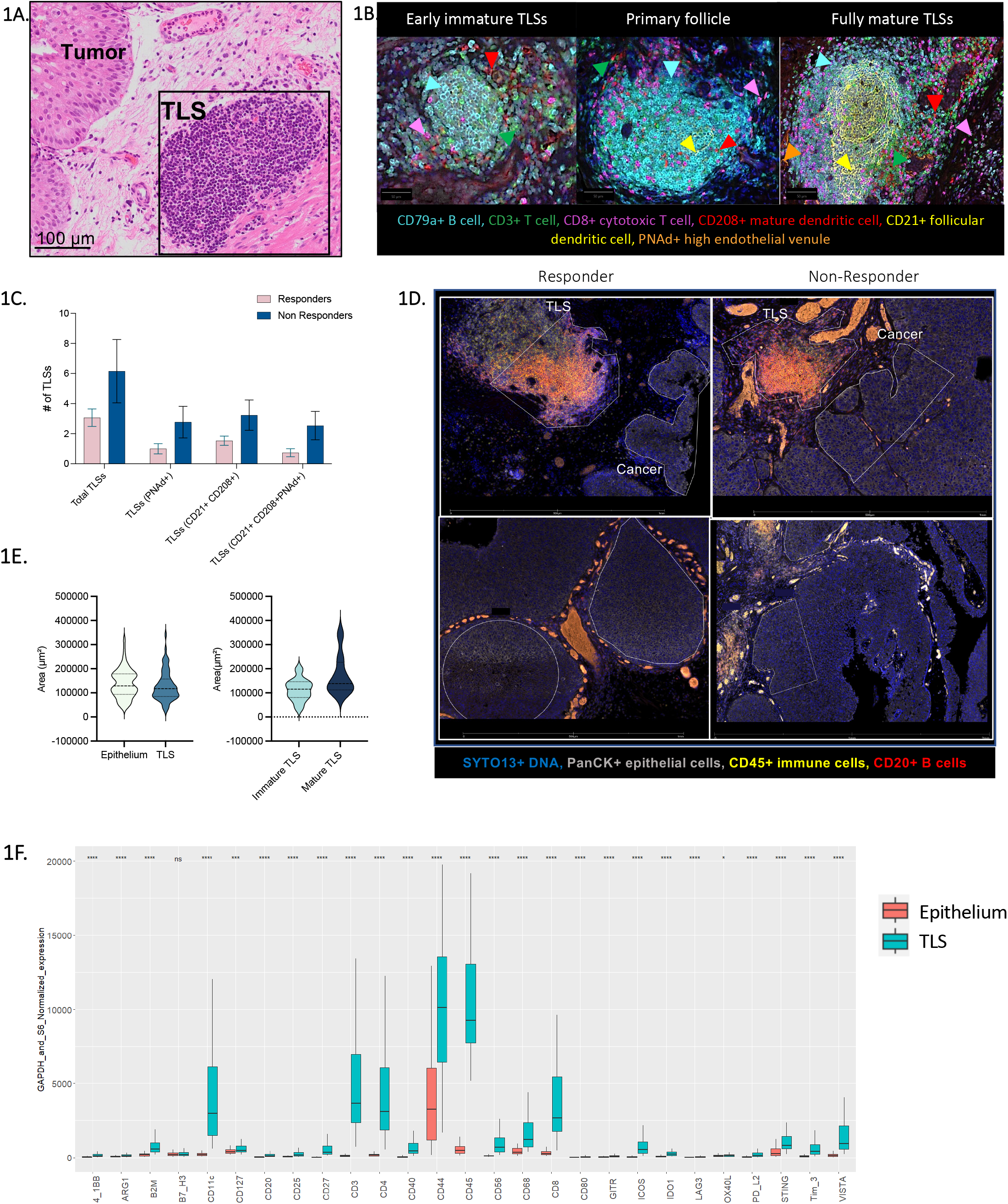
Pre-treatment tumors from BCG non-responders exhibit higher density of tumor adjacent tertiary lymphoid structures. H&E-stained tumor section from a patient classified as BCG non-responder (TLS highlighted in box) **(A)**; Classification of different stages of TLSs (identified by expression of CD21+ and CD208+ follicular dendritic cells and PNAd+ high endothelial venules). Multiplex IF showing different stages of TLSs in whole tumor sections **(B)**. Bar graph showing higher number of TA-TLSs in BCG non-responders (n=13) compared to responders (n=15) **(C)**; Representative images from NanoString GeoMx DSP based staining of 4 whole tumor sections (2 Responders and 2 Non-Responder), showing regions of interest (ROI) selection based on PanCK+/-, CD45+, CD20+ and SYTO 13+ areas/cells **(D)**. Violin plot showing area distribution between TLSs and tumor epithelial compartments in all 12 sections **(E)**. Box plots showing expression of significantly differentially expressed (*p<0.05, ***p<0.001, ****p<0.0001, ns – not significant) proteins of interest between tumor epithelial regions and TLSs **(F)**.

Given the counterintuitive finding on the association of TA-TLSs with early recurrence, we further characterized the expression of 49 immune function related proteins (**Suppl. Table S2)** in TLSs from a subset of 12 specimens (6 in each group of responders and non-responders; **Fig. 1D**) using NanoString GeoMx DSP technology **(Fig. S1)**. The area of mature TLSs was higher compared to all other lymphoid aggregates in the lamina propria (**Fig. 1E**). We did not find any statistically significant differences in the expression levels of proteins associated with immune regulatory functions between TLSs from the responders and non-responders. However, when compared to the tumor epithelial compartment, TLSs from both responders and non-responders exhibited statistically significant higher expression of a) immune regulatory proteins such as CD25, ICOS, TIM3, VISTA, PD-L2, IDO-1; b) antigen presentation associated proteins such as B2M, CD11c, CD40; and c) immune cell phenotype associated proteins, CD45, CD20, CD3, CD4, CD56 and CD68 (**Fig. 1F**). These findings suggest that pre-treatment TLSs in the bladder microenvironment represent chronic carcinogen induced mucosal inflammation and harbor exhausted populations of immune cells at the time of TURBT.

### Chronic exposure to BBN carcinogen increases infiltration of atypical B cells within the bladder microenvironment

To confirm the previously reported age/sex-related shifts in B cell subsets, we first determined the profiles of total B cells and ABCs in the spleen and bone marrow of healthy 12-month-old (reflecting immune profiles of middle-aged humans^18^) C57BL/6 female and male mice. We did not observe any significant differences in the frequency of total B cells (B220+CD19+) from both the spleen and bone marrow of female mice compared to male mice (**Fig. 2A)**; however, splenic ABCs (CD21-/low CD11c+) were significantly higher in 12-month-old female mice compared to their male counterparts (**Fig. 2B – 2D**). Optimization of the B cell-depletion protocol, as per previous reports,^39^ was conducted prior to conducting studies in the carcinogen exposed mice (**Fig. S2B)**.

**Figure 2.**
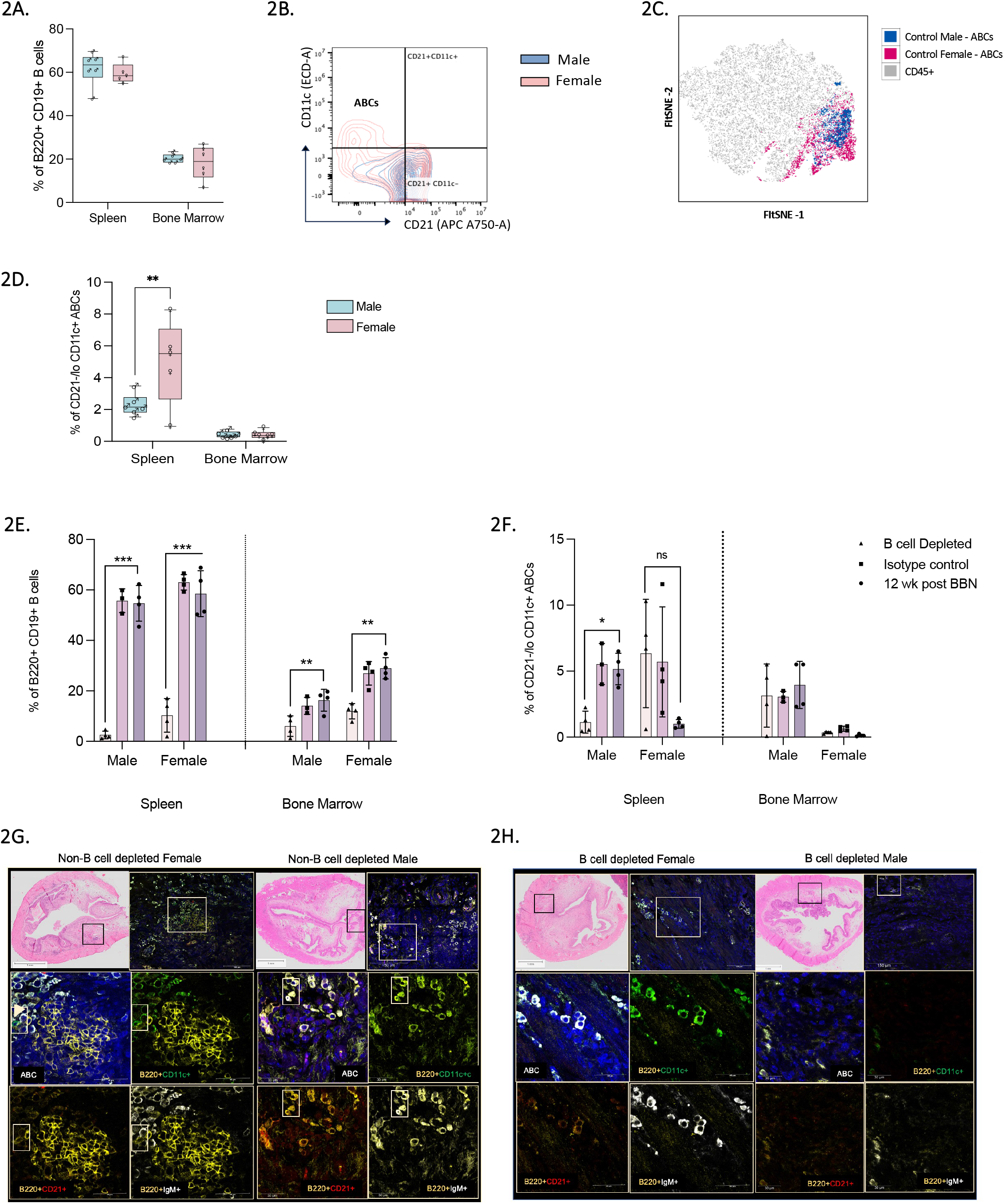
Atypical B cells exhibit sex differences in healthy and carcinogen exposed aging C57BL/6 mice. Comparison of profiles of total B cells (B220+ CD19+) (**A**; n=5-6) and ABCs (CD45+B220+CD19+CD21-/lo CD11c+) (**D**; n=5-6) in spleen and bone marrow of healthy 12-month-old female and male mice. Contour plots and unsupervised Fast interpolation-based t-SNE (FItSNE) plot represents the distribution of ABC population in the total immune (CD45+) cells in healthy male and female mice (**B and C**). FItSNE plot was created using plug-in tool FItSNE and FlowSOM in FlowJo® software. Total B cells and ABCs in B cell depleted, isotype control and untreated female and male mice at 12 weeks post BBN initiation (**E and F**; n=3-4 mice/group). Representative multiplex IF images (n=4/group in each sex) showing differences in bladder infiltrating ABCs at 12 weeks post BBN exposure in female and male mice (**G and H**). Analysis of flow cytometry data was performed using FlowJo® software. All statistical analysis was performed using GraphPad Prism. Ordinary one-way ANOVA with Tukey’s post-hoc test was applied to determine statistical significance in differences (*p<0.05, **p < 0.01, ***p <0.001, ns – not significant). Bar graphs represent mean ± SD.

To determine whether long-term BBN induced genotoxicity and local inflammation leads to increased recruitment of ABCs in the bladder microenvironment, we exposed 12-month-old aging female and male mice to BBN carcinogen for 12 weeks with or without B cell depletion treatment^40^. To specifically determine cause and effect, we first assessed whether B cell depletion during BBN exposure attenuates inflammation in the bladder and delays cancer progression. Results from both local and systemic immune profiling revealed clear sex differences. Female mice displayed an increased frequency of splenic total B cells and ABCs despite depletion **(Fig. 2E and 2F)**. A higher density of ABCs, located with stromal lymphoid aggregates, was observed in the bladder microenvironment of female mice compared to their male counterparts at this time point (**Fig. 2G**). Spatial immune profiling of the bladders collected 1 week post last anti-CD20 injection (12 weeks post BBN initiation), revealed significantly fewer numbers of total B cells and ABCs in the B cell-depleted mice in comparison to the B cell-intact mice (**Fig. 2G and 2H**). Histopathological assessment of whole bladder sections from B cell-depleted and B cell intact mice, revealed a benign or close to normal histology of the bladder urothelium in the B cell-depleted female mice **(Fig. S3A**). Reactive atypia/dysplasia was present in the majority of BBN exposed female and male mice that did not undergo any depletion. Importantly this effect was captured at this time point only in the female mice. These findings suggest increased recruitment (or differentiation) of B cells to the bladder microenvironment due to chronic carcinogen induced mucosal inflammation.

### BBN and BCG induced expansion of splenic ABCs and bladder-associated TLS formation is age and sex-dependent

To determine the role of B cells in treatment response, we treated BBN exposed mice with three weekly intravesical instillations of BCG (**Fig. 3A**). Compared to healthy bladders (pre-BBN; **Fig. 3B**), on hematoxylin and eosin (H&E) staining of bladder sections we observed increased infiltration of immune cells and TLSs at 7 weeks post BBN initiation in the bladder lamina propria of female mice (**Fig. 3C**) in contrast to their male counterparts (**Fig. S3D**). Following the 1^st^ dose of BCG, we observed recovery of the urothelium to its benign morphological states (**Fig. 3D**), in contrast to the saline treated mice where dysplasia/CIS like features were still present at this time point (**Fig. S3B**). However, at 1-, 6-, and 8-weeks post completion of all 3 doses of BCG, we observed urothelial dysplasia and progressive focal to widespread CIS, invasive carcinoma-like changes, or papillary tumors despite removal of BBN exposure at the initiation of BCG treatment in female mice (**Fig**. **3E-3G**). Repeated BCG treatments also led to the formation of TLSs, as confirmed by mIF staining, in contrast to the bladders from saline treated mice where tumor growth was observed with no evidence of TLS formation, (**Fig. S3D and S3E**). An increase in splenic ABCs at 1 week post 1^st^ BCG, compared to the pre-BCG levels, was observed in both female and male mice (**Fig. 3H and 3I**). However, the magnitude of increase was significantly higher in female mice (**Fig. 3I**). A significant decline in the proportion of splenic ABCs post 3^rd^ BCG in both sexes suggests its potential trafficking to the bladder mucosa **(Fig. 3I)**. Additionally, histopathological evaluation **(Fig. 3E)** and mIF **(Fig. 3J and 3K)** of the bladder sections post 3^rd^ BCG revealed presence of ABCs and lymphoid aggregates, particularly prominent in female bladder sections. Furthermore, higher proportion of splenic ABCs after B cell depletion is indicative of accelerated expansion of ABCs potentially due to repeated weekly BCG treatment with simultaneous BBN exposure (**Fig. 3I**). Indeed, removal of BBN exposure at the time of treatment initiation reduced the frequency of splenic ABCs and CD21+ CD11c+ B cells in female mice at one week after the 3^rd^ BCG instillation (**Fig. S7)**. Overall, these findings suggested a combination of carcinogen and BCG treatment induced expansion of ABCs that occurs at a higher magnitude in females. Disease progression post repeated BCG instillations potentially reflects an exhausted immune system.

**Figure 3.**
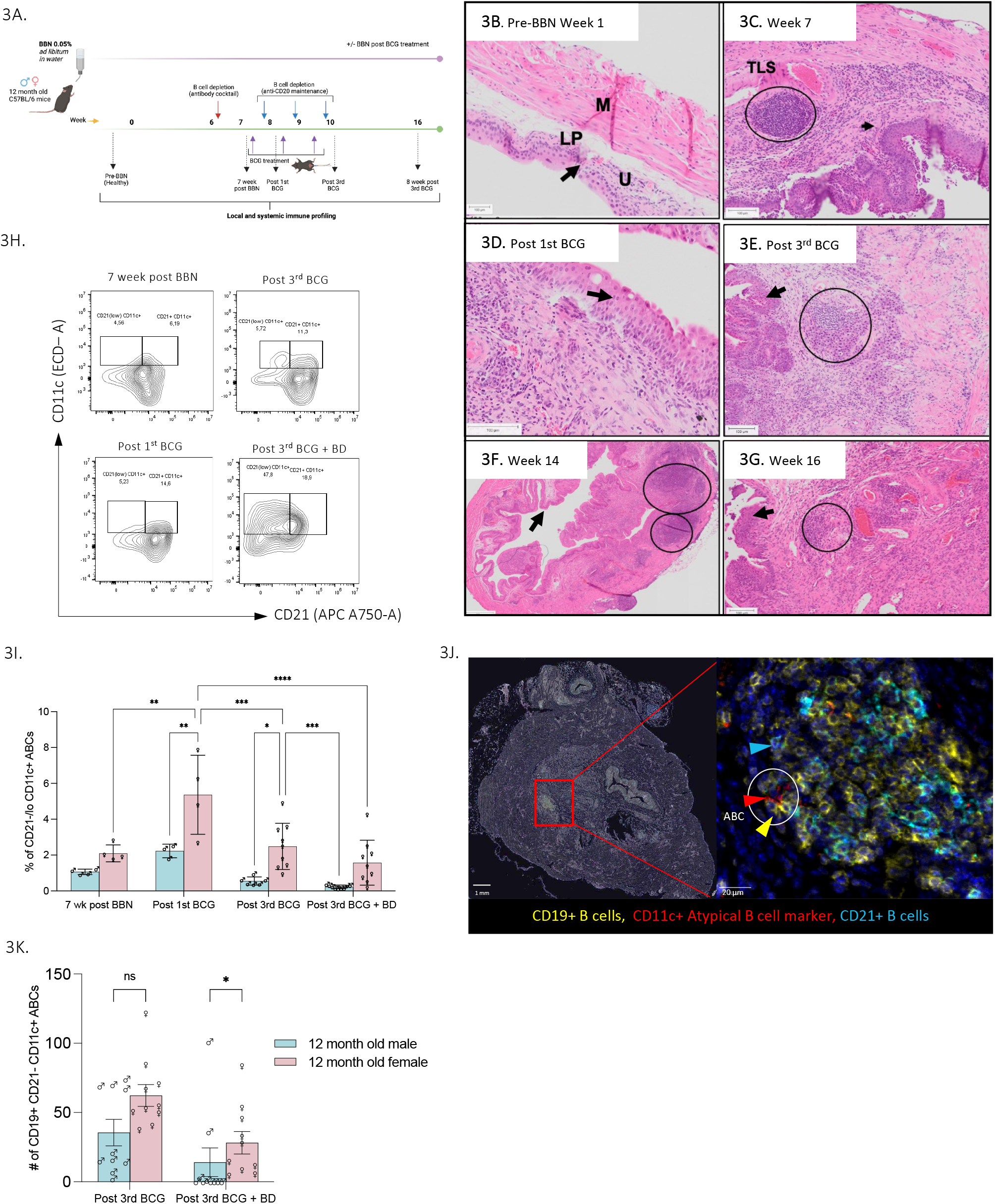
Repeated weekly BCG treatment in aging mice promotes expansion of ABCs and disease progression. Schematic diagram showing experimental approach for intravesical BCG treatment (3 weekly doses) with or without B cell depletion in 12-month-old female and male mice **(A)**. H&E-stained sections showing histological changes in the urothelium and bladder microenvironment at various time points pre-BBN **(B)**, post 7-week BBN **(C)**, one week post 1^st^ BCG **(D)**, one week post 3^rd^ BCG **(E)**, and 6 and 8-weeks post 3^rd^ BCG **(F&G)**. Circles indicate lymphoid aggregates/TLSs. Black arrows indicate the urothelium of bladder sections. Scale bar in H&E-stained images – 100 μm. Representative contour plots for identification of splenic ABC population at different time points post BBN and BCG treatment. Contour plots are illustrative of frequency of CD19+ cells and may show disparity in density in comparison to bar graph as shown in B cell depleted and post 3^rd^ BCG treatment group **(H)**. Bar graph represents differences in the frequency of ABCs (CD19+ CD21-/low CD11c+) post BBN and BCG treatment at different time points for both male (n=3-9) and female mice (n=4-9) **(I)**. Analysis of flow cytometry data was performed using FlowJo® software. All statistical analysis was performed using GraphPad Prism. Ordinary two-way ANOVA with Tukey’s post-hoc test was applied to determine statistical significance in differences (*p<0.05, **p < 0.01, ***p <0.001, ****p<0.0001). Multiplex IF-stained whole bladder sections showing ABCs (CD19+CD21-CD11c+) in 1 week post 3^rd^ BCG treated 12-month-old female mice **(J)**. Representative of 8-10 regions per section (n=5/6 mice in each group). Images at 200x magnification. Bar graph represents number of ABCs (CD19+CD21-CD11c+) detected in 10 annotations per treatment group of the bladder microenvironment **(K)**. Cell detection analysis and classifier to identify different subsets was performed using QuPath software. All statistical analysis was performed using GraphPad Prism. Mann-Whitney test was applied to determine statistical significance in differences (*p<0.05, ns-not significant). Data presented as mean ± SD.

### Sex chromosome associated factors influence B cell response following treatment with BCG

The X chromosome harbors multiple B cell function associated genes such as *CD40L, CXCR3, TLR7, IL9R, IL13RA1/2,* and *BTK*^41^. Furthermore, *TLR7* (known to escape X inactivation) and *IL13RA* are also known to play key roles in the expansion of ABCs. We thus addressed the fundamental question of whether sex-associated ABC expansion in BBN-exposed and BCG-treated mice is influenced by gonadal hormones, sex chromosome complement, or a combination of both factors. We employed 12-month-old aging FCG mice to investigate these interactions (**Fig. 4A**). While no significant differences in the total splenic B cells (B220+CD19+) were observed between the four genotypes; XXF (chromosomal and gonadal females), XYF (gonadal female), XXM (chromosomal and gonadal males) and XYM (gonadal males) of the FCG mice (**Fig. S2C**), the XXF mice exhibited significantly higher ABCs compared to all other genotypes (**Fig. 4B**). We then investigated the sex associated expansion of ABCs in BBN-exposed BCG-treated FCG mice. Splenic ABCs significantly decreased post 3^rd^ BCG dose in the XXF mice compared to the BBN exposed untreated, and saline groups (**Fig. 4B and Suppl. Table S8**). This shift was accompanied by a simultaneous increase in the bone marrow frequency of ABCs at this time point, with a significantly higher increase in XXF mice compared to the XYF, XYM, and XXM mice (**Fig. 4C**). The BBN and BCG associated significant shifts were not observed within any other genotype of the FCG mice indicating that ABC expansion is a plausible combined effect of both X chromosome associated and hormonal factors. Histopathological evaluation revealed the presence of a higher density of lymphoid aggregates in the bladder lamina propria of XXF mice at this time point compared to XYF mice (**Fig. 4D)**. Similarly, H&E staining and mIF revealed an overall higher immune cell infiltration in the bladders of XXM mice compared to the XYM mice (**Fig 4D and Fig. S2D**). These findings confirmed a biological sex and age-associated expansion of ABCs in response to BCG treatment.

**Figure 4.**
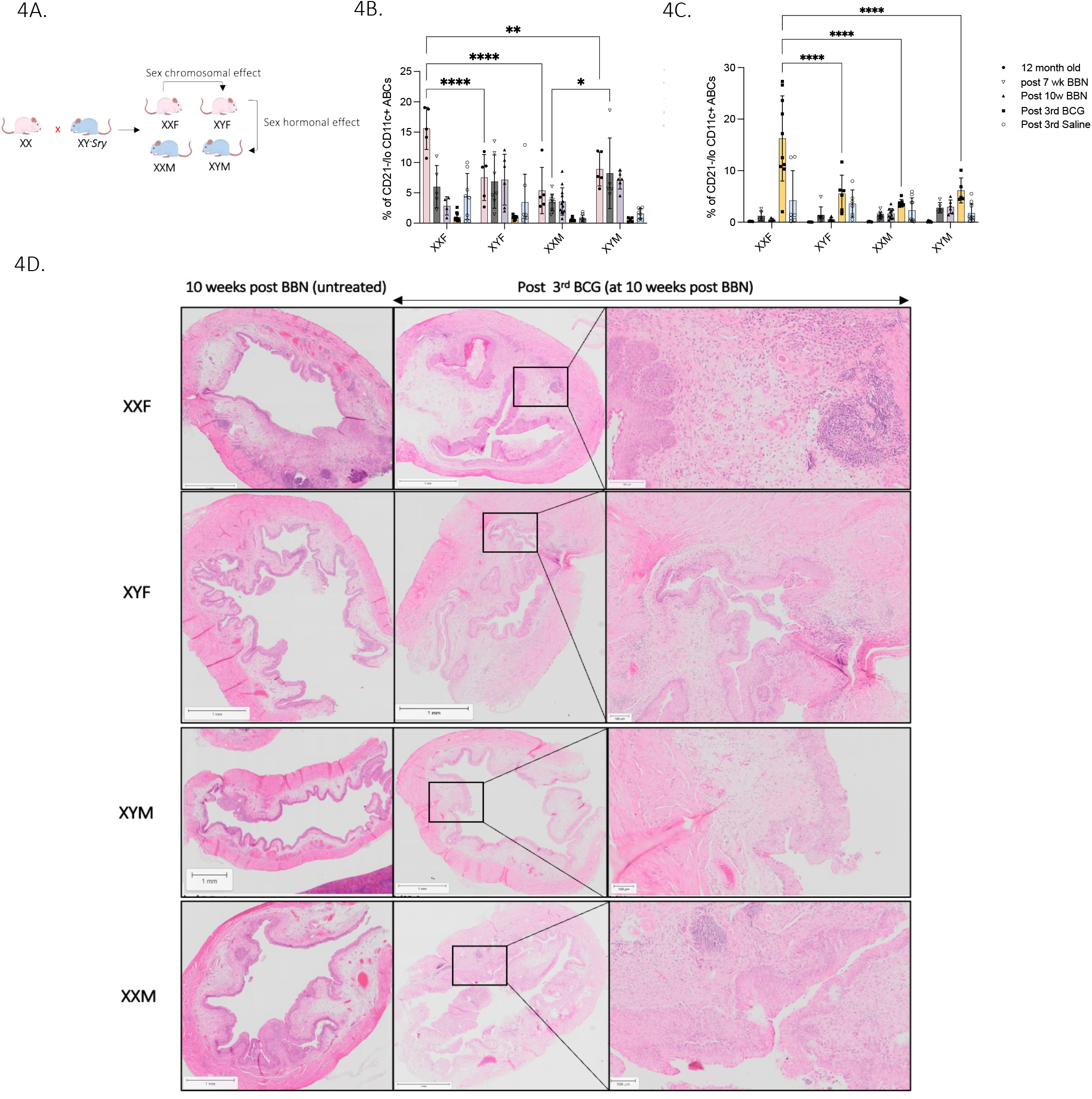
Expansion of ABCs is influenced by hormonal and sex chromosome associated factors. Schematic representation of four core genotype (FCG) mouse model used to study the hormonal and sex chromosomal influence on response to BCG treatment in BBN exposed mice **(A)**. Bar graph showing splenic **(B)** and bone marrow **(C)** CD45+B220+CD19+CD21-/low CD11c+ ABCs in healthy 12-month-old FCG mice; at 10-weeks post BBN and at 1 week post 3^rd^ BCG instillation in BBN exposed mice (BCG treatment initiated at week 7 post BBN initiation). Immune cell frequency was determined using multiparametric flow cytometry. H&E-stained bladder sections from all four genotypes of FCG mice showing differences in bladder microenvironment and overall immune cell infiltration under untreated and BCG treated BBN exposed conditions **(D)**. GraphPad software was used to determine statistically significant differences in profiles across the different genotypes using two-way ANOVA with Bonferroni’s test. (*p<0.05, **p<0.001, ****p<0.0001). Statistical compa_7_rison (p value) among different groups shown in **Suppl. Table S8**. Data shown as mean ± SD. N=3-9 per group. Scale bar in H&E-stained images – 100 μm.

To further confirm the response to ABC phenotype inducing stimuli, we treated splenic total B cells (**Fig. 5A and 5B)**, isolated from BBN exposed 12-month-old female and male mice, to the known ABC inducing cytokines, IFN-γ and IL-21 in the presence or absence of BCG. A significant increase in the frequency of ABCs (CD21-CD11c+) in IL-21 treated B cells isolated from female mice was confirmed *via* multiparametric flow cytometry at 18 hours post-treatment (**Fig. 5C; Fig. S4B)**. Notably, B cells isolated from male mice did not show a similar increase in response to IL-21 treatment (**Fig. 5C)**. However, CD11c+ B cells from male mice exhibited higher proportion of CD21low/+ cells compared to the female mice (**Fig. S4C and S4D)**. The higher expression of CD21 in B cells isolated from male mice potentially suggests a delayed expansion of ABCs following treatment with different ABC phenotype-inducing stimuli. Furthermore, stimulation with a TLR7 agonist (Imiquimod) significantly increased differentiation of splenic B cells, from both female and male mice to ABCs (**Fig. S4E**). Gene expression analysis, using quantitative real time (qRT)-PCR, revealed significant alterations in the expression of ABC associated genes *Fcrl5, Tbx21, and Itgax* at 6 hours post-stimulation (**Fig. 5D-5F).** Treatment of splenic B cells from male mice with IL-21 alone led to a significant increase in the expression *of Itgax* (**Fig. 5D**), indicative of higher ABC differentiation. BCG infection elevated *Itgax* expression in B cells from female mice only when added in combination with IFN-*γ* (**Fig. 5D**). Treatment with IFN-*γ* alone led to similar increase in expression of *Tbx21*, only in B cells from females (**Fig. 5E**). However, in the presence of both BCG and IFN-*γ*, increase in *Tbx21* expression was similar in B cells from both females and males (**Fig. 5E**). In contrast, the combination of BCG infection with IL-21 and IFN-*γ* treatment enhanced *Fcrl5* expression in male splenic B cells (**Fig.5F**). These findings indicate that BCG infection drives a higher differentiation of ABCs in B cells isolated from female mice. Similarly, multiplex cytokine analysis revealed significantly higher levels of GM-CSF, IL-10, IL-6, TNF-*α*, and MCP-1 in the supernatants collected at 18 hours post-treatment of B cells from female mice compared to those from males (**Fig. 5G-5K**). Internalization of BCG by B cells was confirmed *via* immunofluorescence assay using an anti-LAM antibody (**Fig. S4F**). These findings underscore the significance of considering sex-specific responses in B cell differentiation to ABCs following treatment with IFN-*γ* and IL-21 in the presence or absence of BCG bacteria.

**Figure 5.**
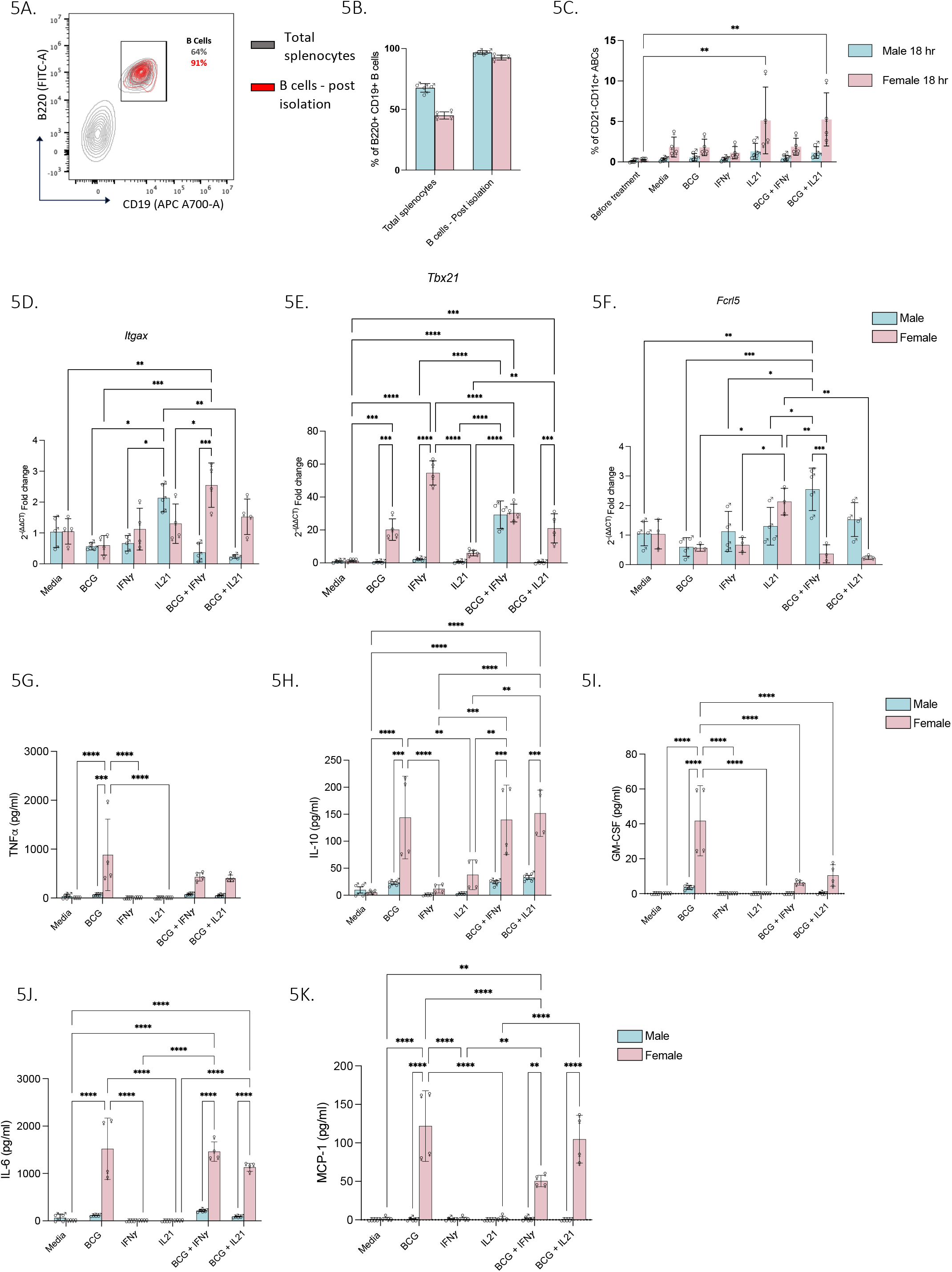
BCG infection and exposure of B cells to IFN-*γ* and IL-21 increases differentiation of B cells to ABCs in a sex differential manner. Representative contour plot for identification of B220+ CD19+ total splenic B cells before and after enrichment **(A)**. Bar graph illustrates the purity of B cells (93.4 ± 2.2%) after isolation from 12-month-old male and female mice exposed to 7 weeks of BBN **(B)**. Bar graph showing increase in ABCs (CD19+ CD21- CD11c+) after 18 h of incubation with IL-21 with or without BCG infection **(C)**. Bar graph represents the *Ubc* house keeping gene-normalized 2^-ΔΔCt^ fold change expression levels of ABC-associated genes *Itgax* (A)*, Tbx21* (B), and *Fcrl5* (C) genes in splenic B cells of BBN exposed male and female mice, at 6 h post treatment with IFN-*γ*, IL-21 in the presence or absence of BCG bacteria. Bar graphs showing significantly increased levels of TNF-*α* **(D)**, IL-10 **(E)**, GM-CSF **(F)**, IL-6 **(G)** and MCP-1 **(H)** in the supernatants collected at 18 hours post incubation of B cells from female mice compared to those from male mice. B cells were treated with IFN-*γ*, IL-21 with and without infection with BCG and subjected to multiplex cytokine profiling. GraphPad software was used to determine statistically significant differences in profiles across the different genotypes using two-way ANOVA with Tukey’s post hoc test. (*p<0.05, **p<0.001, ****p<0.0001). N=3-4 per group. Data shown as mean ± SD.

### B cell depletion during BCG treatment leads to enhanced recovery of the urothelium

Our findings showing ABC expansion during disease progression prompted us to further investigate whether B cell depletion alters the response to BCG immunotherapy in mice with chronic exposure to BBN carcinogen. We found that depletion of B cells prior to and during BCG treatment led to the return of urothelium to its benign morphological state in female mice (**Fig. 6A**). B cell depletion also led to reduced inflammation in the lamina propria of female mice (**Fig. 6A**) compared to male mice (**Fig. 6C**). In contrast, bladders from BCG-treated B cell intact female (**Fig. 6B)** and male (**Fig. 6D)** mice showed dysplasia and focal CIS like changes at 1 week following completion of treatment with 3 doses of BCG. TLS formation was rare or absent in the BCG + B-cell-depleted mice (**Fig. 6E**) compared to B cell intact female mice (**Fig. 6F**). Exposure to BBN carcinogen was maintained throughout the course of BCG treatment and B cell depletion to mimic smoking behavior during BCG treatment. Indeed, removal of the BBN carcinogen at the time of BCG treatment initiation in combination with B cell depletion revealed decreased invasion into the sub-mucosal region (**Fig. S3C**).

**Figure. 6.**
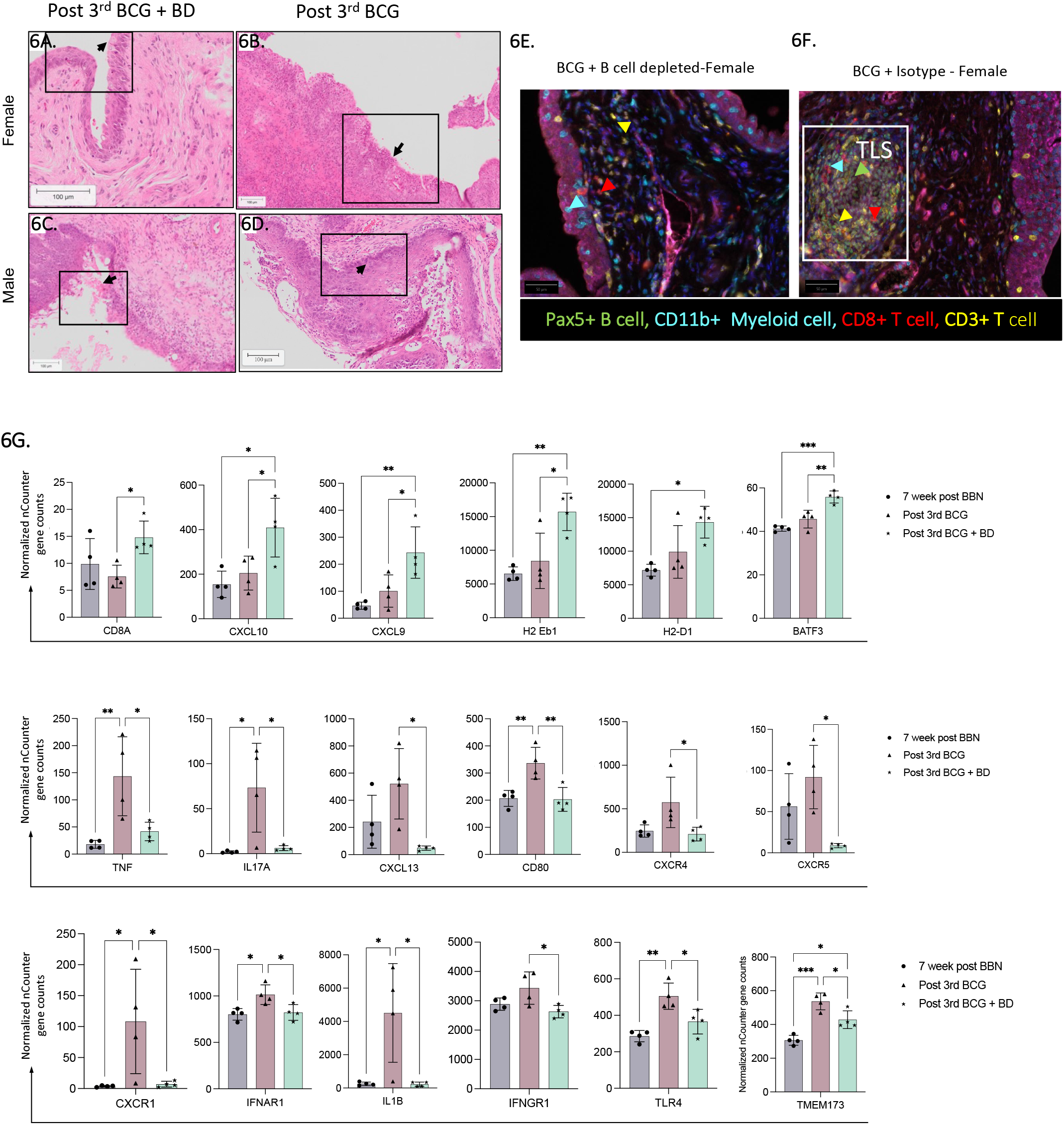
Effect of transient B cell depletion during BCG treatment on urothelium of BBN exposed female and male mice. H&E-stained whole bladder section showing benign/normal histology of urothelium (shown by arrow) post 3^rd^ BCG + B cell depletion in female **(A)** and focal hyperplasia in male **(C)** mice compared to dysplasia and CIS post 3^rd^ BCG treatment in female **(B)** and male **(D)** mice. Multiplex IF stained whole bladder section showing dysplasia and TLS formation (white box) post 3^rd^ BCG instillation in BCG+ isotype treated (F) compared to the BCG+ B cell depleted female mice **(E)**. Representative of 8-10 regions/section, 200x magnification (n=5/6 mice/group). Scale bar in H&E-stained images– 100 µm. Bar graph illustrating the mean ± SD of normalized nCounter gene counts (as indicated), analyzed using NanoString nSolver software. Changes in the expression profiles of selected set of genes in fresh frozen bladders from female mice at 7-weeks post BBN exposure, and one week post 3rd BCG with or without B cell depletion **(G)**. Statistical analysis was conducted using GraphPad software with a one-way ANOVA test to identify significant differences in expression profiles across the various genotypes (*p<0.05, **p<0.001). N=4 per group.

Given the pronounced effect of depletion, specifically on the urothelium of female mice, we examined the profiles of genes involved in immune cell recruitment and TLS formation using a custom NanoString gene panel. The BCG-treated B cell-depleted group demonstrated an increased expression of genes associated with an active effector immune state and antigen processing, such as *Cd8A, Cxcl10, Cxcl9, H2Eb1, H2D1, H2K1, Stat1 and Batf3* (**Fig. 5G; Fig. S5)**. On the other hand, bladders from mice that underwent repeated BCG treatment without B cell depletion showed increased expression of genes associated with the expansion of ABCs and TLS formation, such as *Tnf, Il6, Il17A, Cxcl13, Cd80, Cxcr4, Cxcr5, Cxcr1, Il1b, Ilr1, Tlr4, E2-2, Cd39, and Tmem173* (**Fig 5G; Fig. S5**). These findings suggest that repeated weekly instillations of BCG lead to the systemic expansion and trafficking of ABCs to the bladder microenvironment accompanied by TLS formation. A combination effect of BBN carcinogen and BCG treatment induced sustained inflammation may cause systemic expansion of exhausted immune cells during the contraction phase thus augmenting disease progression.

### B cell depletion during BCG treatment alters the frequencies of helper and cytotoxic T cell subsets

Given that BCG primarily induces T helper type 1 (Th1) responses *via* activation of CD4+ T helper cells, we explored the influence of B cell depletion on the profiles of T cell subsets and whether this occurs in a sex differential manner. We did not observe any significant sex differences in splenic total CD3+ T cell frequency at both early and late (post 3^rd^ BCG) time points (**Fig. 7A and 7B**). Interestingly, the frequency of total CD3+ T cells in the bone marrow was significantly higher in female mice post 3^rd^ BCG compared to male mice (**Fig. 7B**). Notably, in the B cell depleted group there was a significant increase in total splenic T cell frequency compared to the B cell intact group in both sexes at this time point (**Fig. 7B**). Splenic immune profiles of saline treated mice reflected catheterization induced sterile inflammation (data not shown), an effect that is well established in both humans and mice^42,43^. A BCG specific immune response was revealed by the changes in the urothelium (**Fig. S5A**) and the T cell profiles in the bone marrow (**Fig. 7B**) where a significant increase in CD3+ T cell proportions was seen only in the female mice that received 3 doses of BCG with or without B cell depletion. A decline in splenic CD4+ T helper cells and CD8+ T cells following all 3 BCG instillations indicated potential recruitment to the bladder (**Fig. 7C - 7E**). Presence of high CD8+ T cells was confirmed by mIF of bladders collected at the same time point (**Fig. S5A**). Overall, these findings confirm that in response to the administration of BCG *via* intravesical (mucosal) route, the host elicits BCG specific T helper cell associated response that augments a cancer cell specific cytotoxic immune response.

**Figure 7.**
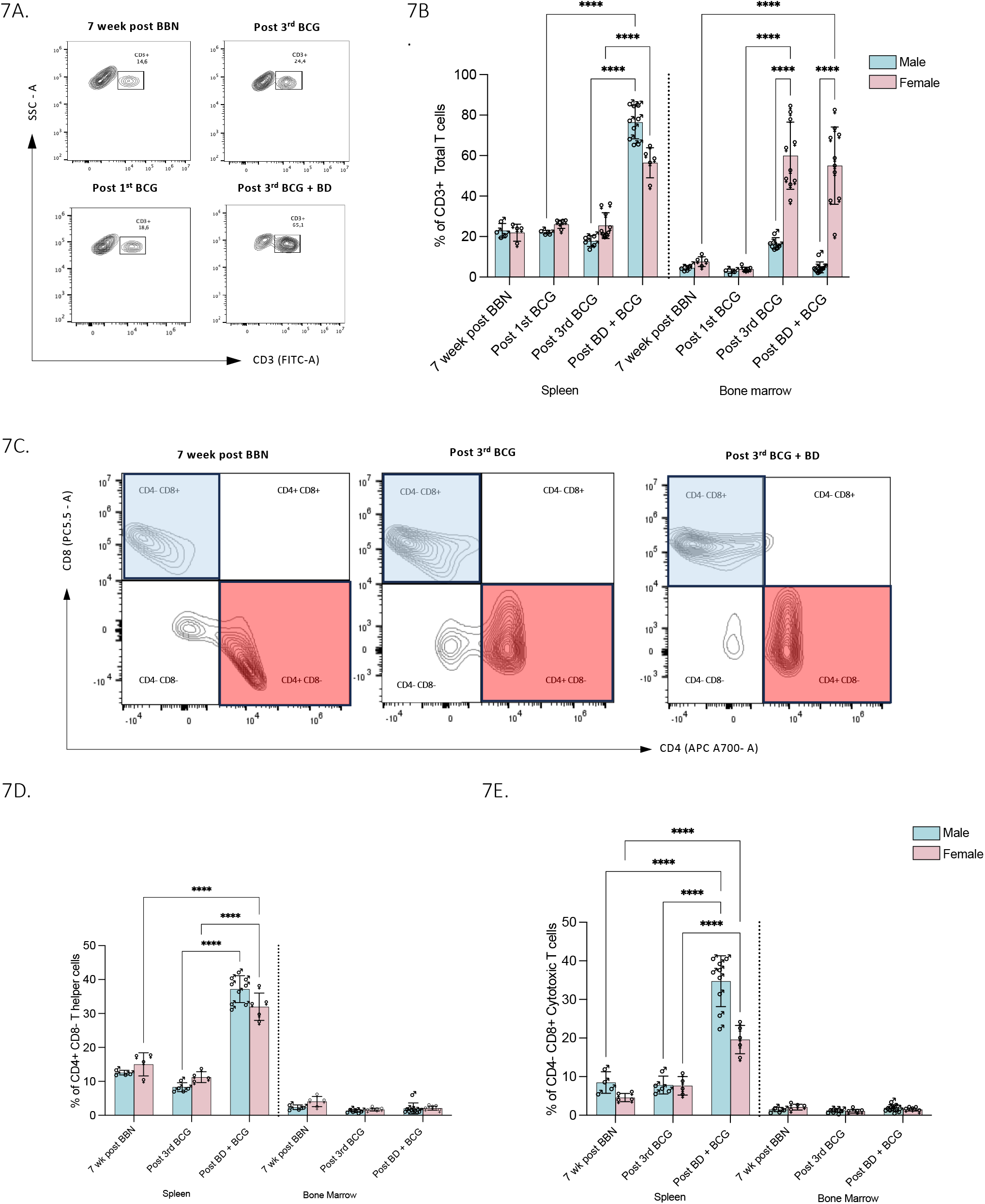
Depletion of B cells during BCG treatment alters systemic T cell profiles in BBN exposed female and male mice. Representative flow cytometry contour plots demonstrating differences in splenic CD45+CD3+ total T cell population in different treatment groups **(A)**. Bar graph represents frequency of total CD3+ T cells in the spleen and bone marrow of 12-month-old male and female C57BL/6 mice at 7-weeks post BBN; one week post 1^st^ BCG; one week post 3^rd^ BCG with or without B cell depletion **(B)**. Representative flow cytometry contour plots demonstrating differences in CD4+ CD8- and CD4- CD8+ T cell subsets in different treatment groups **(C)**. Bar graph illustrating the variations in the frequency of CD4+ CD8- T helper cells **(D)** and CD4-CD8+ cytotoxic T cells (E) in the spleen and bone marrow of 12-month-old aging male and female mice at 7-weeks post BBN; one week post 3^rd^ BCG with or without B cell depletion. Analysis of flow cytometry data was performed using FlowJo® software. All statistical analysis was done in GraphPad Prism software using two-way ANOVA with Bonferroni’s test depicted statistical significance. (****p < 0.0001). N=3-9 per group. Data presented as mean ± SD.

### B cell depletion during BCG treatment alters systemic and bladder infiltrating myeloid cell functional states

Bladder resident and recruited populations of myeloid cells are key to BCG response. In this study we observed that BCG treatment led to significant alterations in the profiles of CD11b+ myeloid cells in the spleen and bone marrow of both female and male mice. Overall, female mice depicted a higher proportion of myeloid cells in the spleen compared to males. A small but significant increase in splenic total CD11b+ myeloid cells, at 1 week post 1^st^ BCG dose, was only observed in the female mice compared to the male mice (**Fig. 8A and 8B).** Following completion of treatment with all 3 doses of BCG, in both female and male mice, there was a significant decline in bone marrow CD11b+ cells in contrast to mice undergoing B cell depletion during BCG (**Fig. 8B**). A significant increase in splenic PD-L1 immune checkpoint expressing myeloid cells was observed only in female mice following the 1^st^ BCG dose (**Fig. 8C**). The proportion of PD-L1 expressing myeloid cells was significantly higher following the 3^rd^ BCG dose in the spleen and bone marrow of both female and male mice that underwent B cell depletion during BCG treatment (**Fig. 8C and 8D)**. This was also confirmed by spatial immune profiling of bladders from the same mice where increased infiltration of macrophages was observed (**Fig. 8E-8G).** Spatial immune profiling of bladders revealed that the BCG + isotype treated female mice had increased infiltration of PD- L1+ cells than the B cell-depleted group (**Fig. 8E-8G**). These findings suggest that the combination of BBN exposure and repeated weekly BCG treatment induces sustained local inflammation, which could potentially accelerate the exhaustion of systemic and bladder local antigen-presenting cells (APCs).

**Figure 8.**
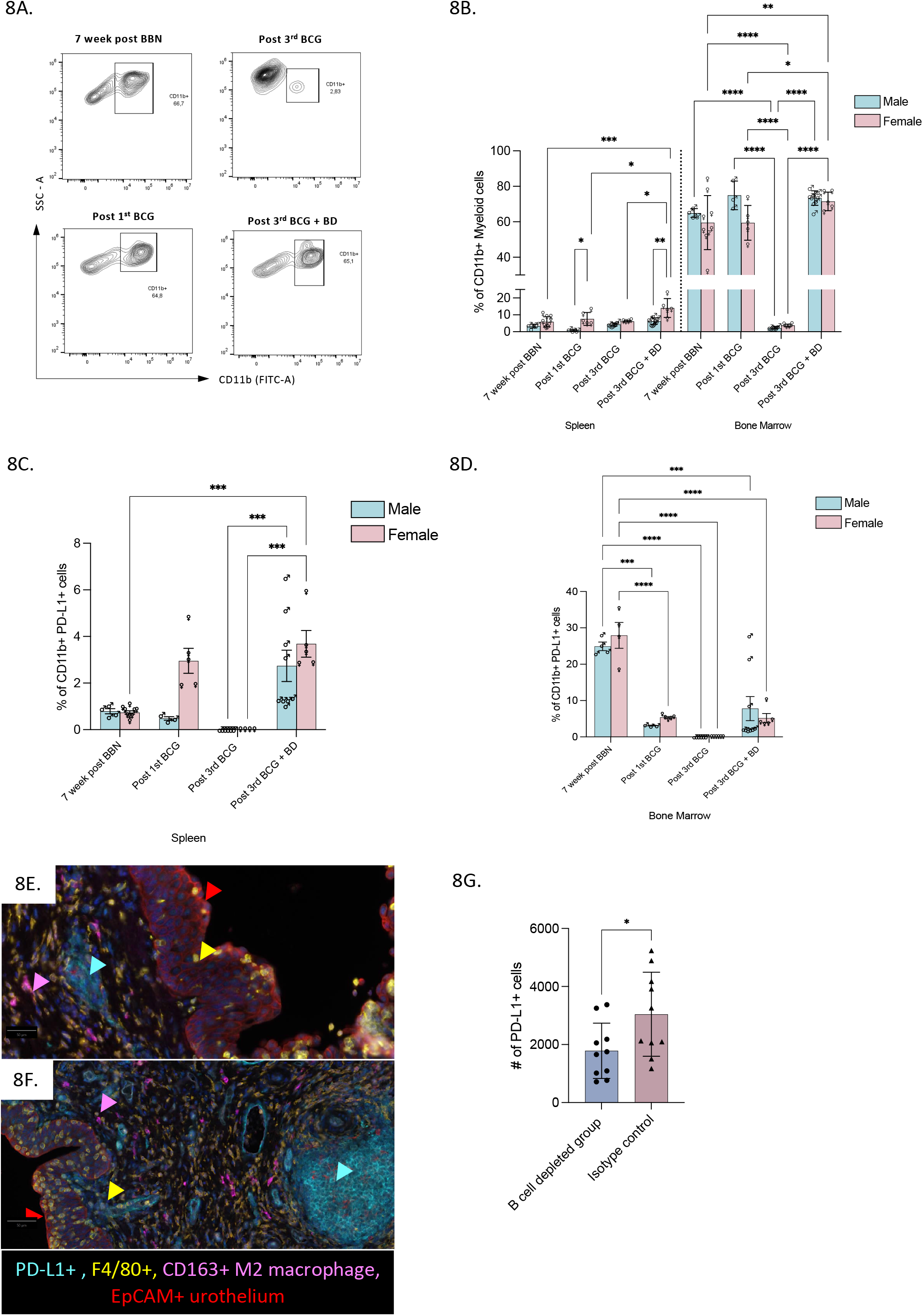
Repeated BCG treatment increases PD-L1 expression on splenic myeloid cells and in the bladder microenvironment. Representative flow cytometry contour plots demonstrating differences in splenic CD45+ CD11b+ total myeloid cell population in different treatment groups **(A)**. Bar graph represents profile of total CD11b+ myeloid cells in the spleen and bone marrow of 12-month-old male and female C57BL/6 mice at 7-weeks post BBN; one week post 1^st^ BCG; one week post 3^rd^ BCG with or without B cell depletion **(B)**. Bar graph represents differences in the profile of CD11b+ PD-L1+ cells in spleen **(C)** and bone marrow **(D)** of 12-month-old aging female and male mice at 7-weeks post BBN; one week post 1^st^ BCG; one week post 3^rd^ BCG with or without B cell depletion. Analysis of flow cytometry data was performed using FlowJo® software. All statistical analysis done in GraphPad Prism software (v9.5.1) using two-way ANOVA with Bonferroni’s test depicted statistical significance. (*p<0.05, **p<0.01,***p < 0.001,****p < 0.0001). Multiplex IF-stained whole bladder sections showing PD-L1+ cells in BCG + B cell depleted **(E)** and BCG + isotype antibody treated mice **(F)** at 1 week post 3^rd^ BCG instillation. Representative of 8-10 regions per section. Images at 200x magnification. Bar graph (mean ± SD) represents PD- L1+ detections in 10 annotations per treatment group of the bladder section **(G)**. Cell detection analysis and classifier to identify PD-L1+ expression was performed using QuPath software. All statistical analysis was performed using GraphPad Prism. Mann-Whitney test was applied to determine statistical significance in differences (*p<0.05). N=3-10 per group. Data presented as mean ± SD.

### B cell depletion during BCG treatment of BBN exposed mice leads to elevated levels of plasma Th1 and Th2 proinflammatory cytokines and antibodies in female mice

Multiplex cytokine profiling of plasma collected following the completion of 3 doses of BCG revealed significant differences in pro-inflammatory and immune cell activating cytokines key to mediating Th1 and Th2 responses. In female mice undergoing B cell depletion during BCG treatment, plasma levels of the Th1 cytokines; IL-2, IFN-γ, GM-CSF, MIP-1α and MIP-1β were significantly higher compared to mice that did not undergo B cell depletion during BCG treatment. Similarly, significantly increased Th2 cytokines such as IL-9 and IL-13 were observed in mice subjected to B cell depletion during BCG treatment (**Fig. 9**). Healthy female mice displayed higher plasma levels of IgM, IgG2a and IgG2b, compared to the age-matched male mice (**Fig. S10B, S10C and S10D**). However, the levels of IgG2a declined at 7-week post BBN and 1 week post 3^rd^ BCG in female mice **(Fig. S10C**). Notably, in the female mice, the levels of IgG2b increased from their pre-BCG levels reaching up to similar levels to those in age matched healthy controls, despite the depletion of B cells during BCG treatment (**Fig. S10D**). In contrast, IgG2b levels decreased significantly in B cell-depleted male mice (**Fig. S10D**). These findings suggest a critical role of B cell subsets in inhibiting Th1 and Th2 responses induced by repeated mucosal BCG administration in female mice.

**Figure 9.**
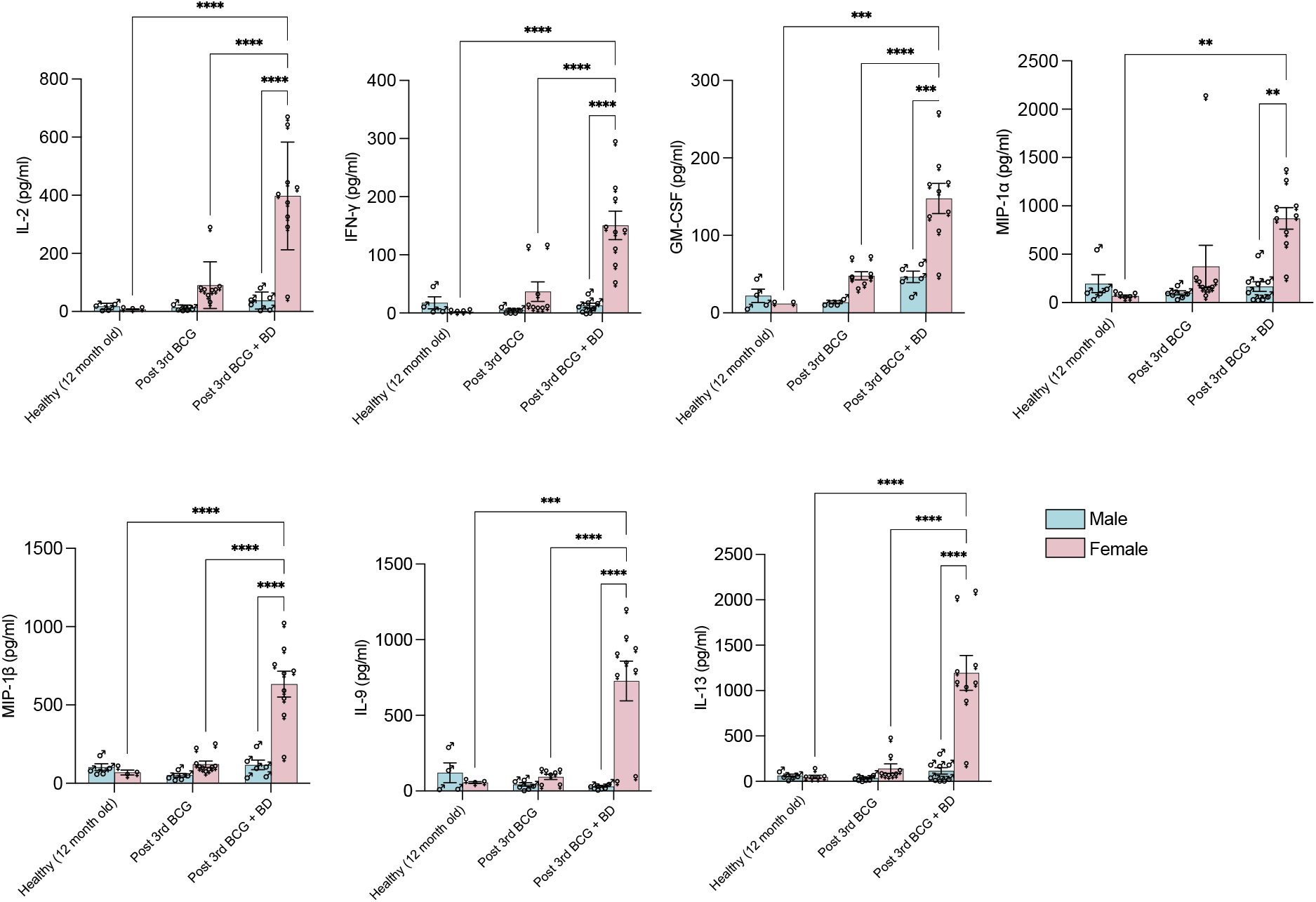
B cell depletion during BCG treatment increases plasma Th1/Th2 cytokine levels. Plasma collected at 1 week post 3^rd^ BCG dose with or without B cell depletion was subjected to multiplex cytokine profiling. Significantly increased levels of Th1 (IL-2, IFN-γ, MIP-1α, MIP-1β), Th2 (IL-9 and IL-13) cytokines and GM-CSF, were observed in BCG+B cell depleted female mice (n=3-4 in each group). All statistical analysis was performed using GraphPad Prism. Statistical significance, *p<0.01, **p<0.001, ***p<0.0001, ****p<0.0001, was determined using two-way ANOVA with Tukey’s post-hoc tests. Data presented as mean ± SD.

## Discussion

B cells and TLSs have emerged as biomarkers of favorable prognosis and response to chemotherapy or immune checkpoint inhibitor therapy in some cancers including muscle-invasive BC (MIBC)^44–46^. Recent reports also show that increased density and proximity of TA-TLSs is associated with improved survival in patients with MIBC who are treated with adjuvant chemotherapy or immune checkpoint inhibitors^47,48^. TA-TLSs are also associated with favorable prognosis for patients with melanoma, renal, gastric, and lung cancers^49,50^. Contrary to this, in colorectal, breast and hepatocellular carcinoma^51^, a negative relationship between TA-TLSs and patient outcomes has been demonstrated^52,53^. Similar to these and our previously reported findings from pre-treatment tumors of patients with NMIBC and MIBC, findings from the current study suggest their negative association with response to BCG^12^. This association may primarily be attributed to the difference in local immunomodulation and downstream responses induced by live BCG bacteria compared to systemically administered immune checkpoint inhibitors or cytotoxic chemotherapy. The discrepancy also highlights the fact that the relationship between TA-TLSs and cancer patient outcome depends on several parameters, including functional states of immune cells within TLSs, persistence of carcinogen, treatment type, age/sex and disease stage. While we observed high expression of immune cell exhaustion associated proteins in chronically evolved pre-BCG TA-TLSs, it is plausible that the induction of acute TLSs post-BCG would be associated with positive clinical outcomes given the ability of BCG to induce the formation of granulomas^54^. However, this association may be reversed depending on the persistence of carcinogen and recruitment of exhausted immune cell populations, from the systemic circulation following repeated exposure to live BCG bacteria (as observed in our *in vivo* studies).

TLSs may also play a pathological role by promoting auto-antibody production and harboring immunosuppressive and dysfunctional immune cell populations due to sustained local inflammation^55^. Indeed, previous reports in NMIBC showed significantly increased levels of circulating IgG antibodies in patients exhibiting early recurrence post BCG^56^. This duality helps to explain our findings as to why TLSs have been strongly associated with both positive and negative outcomes in different cancer types. In this study, mIF based characterization of TLSs did not reveal any significant differences in the density of TLSs in tumors from patients who subsequently responded, compared to those who exhibited early recurrence post BCG. However, the location and maturity of TLSs and proximity to tumor epithelium/invasive margin was indicative of an aggressive tumor behavior accompanied with an exhausted local immune response. Nevertheless, our findings from a cohort of 28 patients (89% male patients) provide novel insights into TLS biology and emphasize the need for further validations in large independent cohorts.

BCG immunotherapy in NMIBC is unique with respect to its mucosal route of administration and dosing schedule that requires repeated intravesical instillations at weekly intervals. Such repeated exposure to BCG antigens, specifically during the induction phase, may cause an expansion of exhausted immune cell populations in the effector memory phase. Such an expansion may occur specifically in BCG unresponsive patients with a history of chronic or persistent carcinogen exposure, smokers, patients with autoimmune conditions or female patients with a history of recurrent urinary tract infections. The recent report by Strandgaard and colleagues^57^ demonstrated T cell exhaustion as a major factor underlying BCG failure. It is plausible that in some patients, such repeated BCG induced immune activation will also lead to increase in exhausted memory B cells and their recruitment to the bladder microenvironment leading to less effective anti-tumor immune responses. This effect will be more pronounced in older patients or those who exhibit higher pre-BCG intra-tumoral density of B cells within TA-TLSs, reflective of their age-related/chronic carcinogen exposure induced profiles. It is tempting to speculate that analogous to this T-bet transcription factor associated fate in both T and B cells, expansion of ABCs (during the effector memory phase) may result from repeated encounter with BCG bacteria in the induction phase, specifically, in unresponsive patients. This effect may potentially also be more pronounced in female patients who generally experience early recurrence and progression. Indeed, our previous report and unpublished findings from an on-going study suggest that both female and male patients with high pre-treatment intra-tumoral B cell density also experience early recurrence post BCG^12^.

Patient age is an independent risk factor associated with poor response to BCG immunotherapy in NMIBC^58^. Immunosenescence associated B cell alterations, governed by hormonal and sex chromosome associated factors, accompany aging in a sex differential manner^23^. Indeed, immune profiling of healthy aging mice confirmed the higher frequency of ABCs in females compared to their male counterparts. Most importantly, using the FCG mouse model, we clearly established that such sex differences in BCG induced systemic immune profiles as well as bladder associated lymphoid aggregate formation, are influenced by both gonadal hormones and X chromosome linked factors. Based on our *in vivo* findings, we reason that BCG internalized by ABCs in the bladder microenvironment/TLSs, activates the TLR9 pathway following the binding of bacterial CpG DNA^35^. Such an amplified TLR9 response could lead to the production of immunosuppressive IL-10, which may further inhibit anti-tumor T cell responses and polarize macrophages to M2-like suppressive states. *In vitro* infection of B cells using BCG bacteria confirmed such an increase in IL-10. Further, significant decrease in PD-L1 expression in bladders B cell-depleted BCG-treated female mice reflects a favorable anti-tumor immune response.

Validation to the central role of B cells, in mediating responses to BCG, is reinforced by noteworthy observations from our study. Notably, the depletion of B cells during BCG treatment reduced inflammation in the urothelium and helped accelerate its recovery despite the presence of carcinogen specifically in female mice. Additionally, the recovery of urothelium was accompanied by significant increases in circulating levels of the cytokines such as IL-2, IFN-γ, IL-9 and IL-13 that favor a Th1 and Th2 response. These cytokines play crucial roles in enhancing the immune system’s ability to fight against cancer cells and facilitate the resolution of inflammation. Furthermore, increased expression of *Cd8A, Cxcl10, Cxcl9, H2-Eb1, H2-d1,* and *Batf3* was observed in bladders from mice undergoing B cell depletion during BCG treatment. A higher expression of *Cd8A* in the depleted group indicated a higher T cell density, which is also supported by our findings from mIF of whole bladder sections from the depleted group. Conversely, the upregulation of *Batf3* suggests either increased DCs or the suppression of T regulatory cells, in the depleted group. On the other hand, the BCG treated group exhibited higher expression of genes linked to ABC expansion and TLS formation, such as *Tnf, Il6, Il17, Cxcl13, Cxcr4, Cxcr5, Cxcr1, Cd80* and *Il1B.* These findings align with our hypothesis that ABC expansion and recruitment following repeated BCG treatment could contribute to an exhaustive bladder microenvironment, potentially leading to early disease recurrence. These gene expression patterns further corroborate the differential immune responses observed between the two treatment groups, emphasizing the pivotal role of B cells in shaping the response to BCG immunotherapy. Despite the depletion of B cells, comparable levels of IgM, IgG2a, and IgG2b between the healthy control and B cell-depleted female mice is suggestive of a memory B cell response triggered by BCG infection given the enhanced rejuvenation of lymphopoiesis in females. It is known that the cytokine IL-21 (inhibits CD8+ T cell cytolytic function), which governs ABC expansion, plays opposite roles compared to those of IL-2 (promotes cytotoxic function of CD8+ T cells)^59^. B cell depletion prior to initiation of BCG treatment may thus ablate such IL-21 mediated suppression of T cell function thus enhancing anti-tumor immunity especially in females or males with high TA-TLSs.

Our study is not without limitations. We did not establish the precise *in vivo* cellular trafficking associated events following BCG. However, our finding that CD3+CD8- T cells were significantly expanded post 3^rd^ BCG in both B cell depleted and non-depleted mice, suggests that BCG bacteria travel to the bone marrow where further cellular priming and activation may occur. While findings from this study demonstrate the critical role of B cells and ABCs in mediating response to BCG, the mechanistic basis of how these cells suppress anti-tumor cytotoxic response remains unclear and needs further investigation. Future investigations could also explore the BCG dose related aspects of differential response to BCG in the context of sex related immune physiology. It is established that B cell associated CD21 (complement receptor 2) binds to autoantigens or immune complexes,^60^ and also interacts with TLR9. Most importantly CD21/CR2 has been shown to be critical for the internalization of BCG bacteria by B cells.^61^ The role of the complement cascade in mediating response to BCG in NMIBC has not been explored and needs further investigation.

Overall, this study not only advances the current understanding of the BCG mode of action in the context of bladder mucosal immunity but also provides the first evidence for a critical role of B cell exhaustion, reflected by higher pre-BCG TA-TLSs and expansion of ABCs, in disease progression and treatment response. While mouse models do not reflect the disease complexity observed in human scenarios, our study considers aging aspects of the immune system, unique aspects of the mucosal immune responses and sex differences *via* the inclusion of mice from both sexes and the FCG mice. Taken together, these novel findings based on human specimens and *in vivo* studies, partially explain the poor outcomes in female patients and in male patients with high intra-tumoral B cells and TA-TLSs in their pre-treatment tumors. Therapeutic targeting of ABC associated markers in combination with optimal sequencing of PD-L1 immune checkpoint blockade could be a novel approach for treatment of patients identified early based on their systemic and bladder associated immune profiles.

## Materials and Methods

### Human NMIBC tumor specimens

All studies involving the use of patient materials were approved by the Human Ethics Review Board at Queen’s University. The cohort comprised of 28 patients (25 males and 3 females) with NMIBC who underwent TURBT surgery at Kingston Health Sciences Centre (KHSC) between March 2016 and November 2017. Formalin-fixed paraffin-embedded (FFPE) tumor specimens were retrieved from the Department of Pathology at Queen’s University. Patient clinical details were retrieved *via* electronic chart review (DJ). All patients in the cohort received adequate induction of BCG immunotherapy (>5 out of 6 doses) following index TURBT surgery. Response to BCG was defined as the time from each patient’s earliest TURBT resection to next malignant diagnosis. Operative notes were reviewed to exclude re-resections as recurrences. Patients who had recurrence-free survival greater than 2 years were classified as BCG-responders (n=15). Patients who had recurrence-free survival less than 1 year were classified as BCG-non-responders (n=13).

### Multiplex immunofluorescence on whole tumor sections

We employed mIF to characterize the organization and maturity of TLSs found in tumors from NMIBC patients. Using our selected archival FFPE samples, triplicate 5 µm sections were cut and mounted at Queen’s Laboratory for Molecular Pathology (QLMP). One section per sample was subjected to H&E staining. Stained sections were subjected to high-resolution scanning (Olympus VS120 Scanner) and the scans were uploaded for viewing on the web-based HALO® image hosting service. Immune cell aggregates, mature TLSs, and other areas of interest were electronically annotated and reviewed for accuracy by a trained pathologist (MX).

An unstained section from each of the samples was stained *via* automated mIF staining at the Molecular and Cellular Immunology Core facility (Deeley Research Centre; Vancouver, British Columbia, Canada). Samples were stained with fluorophore-conjugated primary antibodies against cell surface proteins to identify B cells, T cells, DCs and high endothelial venules (**Suppl. Table S4**). Stained sections were scanned using the Vectra multispectral imaging system and viewed using PhenoChart 1.1.0 Software (Akoya BioSciences®; Marlborough, Massachusetts, USA). Aggregates/TLSs were classified into primary (immature) or secondary (mature) follicle as previously reported.^19^

### Digital spatial profiling of immune pathway proteins using NanoString’s GeoMx platform

We employed NanoString GeoMx™ DSP technology to spatially characterize the protein expression profiles in TLSs and epithelial regions within a subset of NMIBC tumors (6 responders and 6 non-responders). This unique, high-throughput multiplexed technique uses indexing oligonucleotides to quantify the spatially resolved abundance of protein or RNA from FFPE tissue sections. Unstained fresh-cut 5 µm sections from the selected samples were processed on the NanoString GeoMx DSP platform at the Imaging Centre (University of Minnesota, USA). Slide processing was performed as per the manufacturer’s instructions using DNA stain (SYTO 13), pan-Cytokeratin (epithelium), CD45 (immune cells) and CD20 (B cells) antibodies to guide selection of regions of interest. The abundance and localization of 49 immune proteins of interest (Immune Cell Profiling Core, plus the Immuno-Oncology, Drug Target Module, Immune Activation Status Module, and the Immune Cell Typing Module; see **Suppl. Table S2**) was measured. Quality control was performed as per the manufacturer’s instructions and External RNA Controls Consortium in DSP suite (nCounter™ platform) prior to further analysis. Details of the DSP methodology and analysis are provided in the Supplementary data.

### In vivo studies

#### Animal models

All murine experiments were in accordance with the guidelines provided by Queen’s University Animal Care Committee and approved in an animal research protocol. The B6.Cg-Tg(Sry)2Ei Srydl1Rlb/ArnoJ mice were kindly provided by Dr. Manu Rangachari (University of Laval) and also purchased from the Jackson laboratory (stock number 010905, RRID:SCR_002187). Breeding of the FCG mice was conducted at the Queen’s animal care facility to derive the four genotypes (gonadally female mice and male mice with XX and XY chromosomal complement each). The FCG transgenic mouse model (on a C57BL/6 background) allows for the generation of sex-reversed XX males and XY females by breeding mice with a loss-of-function mutation in the Y chromosome-linked testes-determining *Sry* gene with transgenic mice carrying an *Sry* rescue transgene on an autosome (**Fig. 4A**)^62^. This model permits determination of independent effects of gonadal and sex chromosome-specific features on bladder cancer progression associated immunologic profiles. Mice were aged for 12 months at the facility. Wild type C57BL/6 female and male mice (12 months old) were purchased from Charles River Laboratory (Montreal, QC, Canada). All mice were fed sterilized conventional diet and water and maintained under specific pathogen-free environment.

#### BBN carcinogen exposure and depletion of B cells

Female and male C57BL/6 mice (12 months old) and FCG mice (12 months old) were administered 0.05% BBN (Cat# B0938, TCI America, OR, USA) *ad libitum* in drinking water (protected from light) once a week for 7-12 weeks as previously described.^18^ Our previous findings have demonstrated the formation of lymphocyte aggregates, within the lamina propria of bladder, initiating at week 4 post BBN exposure as a result of BBN induced DNA damage^40^. To determine the influence of B cells on disease progression, we initiated B cell depletion starting week 3 post BBN exposure. Briefly, a cocktail of monoclonal antibodies (**Suppl. Table S3**) at 150 μg/mouse: rat anti-mouse CD19, rat anti-mouse B220 and mouse anti-mouse CD22 was injected *via* intraperitoneal (I.P) injection, as described previously^39^. After 48 hours, mice were injected with mouse anti-rat κ secondary antibody at 150 μg/mouse. To maintain the depletion of B cells *in vivo*, mice were injected with anti-CD20 antibody. B cell depletion cocktail injections were repeated 3 times every 10 days to ensure the complete depletion of systemic B cells, further confirmed with flow cytometry results post last injection. To determine the effect of B cell depletion on response to BCG in BBN exposed mice, depletion protocol was initiated 1 week prior to 1st BCG instillation followed by weekly anti-CD20 antibody injections for 3 weeks during BCG treatment. Isotype controls for each antibody (**Suppl. Table S3**) mentioned above were also used.

#### Intravesical BCG in BBN exposed mice

Lyophilized TICE strain of BCG (Merck, Canada) was obtained from the pharmacy at KHSC. Each 50 mg vial contained 1 - 8 x 10^8^ colony forming units (CFU) of BCG, as tested by the manufacturer. BCG was resuspended in sterile saline for all experiments. At 7 weeks post BBN exposure, mice were subjected to intravesical treatment with 3 weekly instillations of BCG (1mg/mouse in 50 μl of saline). BCG was delivered *via* transurethral catheterization with a 24-gauge catheter (3/4”, Cat# 4053, Jelco, Minneapolis, USA) for female mice and PE-10 tubing in male mice under isoflurane anaesthesia. Urethral VASCU-STATT Midi clamps (Cat# 1001-501, Scanlan, Canada) were placed to hold BCG within the bladder for 1 hour. Whole bladders were collected from all treatment groups post 1st (8 weeks post BBN exposure) and 3rd BCG (10 weeks post BBN exposure) instillation, post 1st anti-CD20 treatment after 1st BCG instillation (8 weeks post BBN exposure) and post 3rd anti-CD20 treatment after 3rd BCG (10 weeks post BBN exposure) instillation (**Fig. 3A**) and fixed in 4% paraformaldehyde for histopathological evaluation and spatial immune profiling *via* mIF. Single cell suspensions from bone marrow and spleen were subjected to profiling using multispectral flow cytometry. Blood was collected from the submandibular vein for plasma cytokine profiling.

#### Local and systemic immune profiling

Whole bladders collected from all treatment groups, at treatment time points as indicated in **Fig. 3A**, were sagittaly cut into two halves, fixed in 10% formalin and paraffin-embedded for H&E staining as described in above section. The urothelial changes in mice were classified as follows: reactive atypia, dysplasia, carcinoma in situ (CIS), early invasion and invasion. 5 μm unstained FFPE sections were subjected to spatial immune profiling using mIF staining to determine the infiltration patterns of selected immune markers. Antibodies to identify CD11b+ myeloid cells, CD3+ total T cells, CD8+ cytotoxic T cells, Pax5+ B cells, CD19+ CD21- CD11c+ atypical B cells, PD- L1+ cells (**Suppl. Table S4**) were used in mIF staining at the Molecular and Cellular Immunology Core (MCIC) facility, BC Cancer Agency as per previously reported methods.^18^ Multiplex IF stained images were acquired using the Vectra 3 multichannel imaging system. Images were then imported to Phenochart (11.0, Akoya Biosciences) and Inform Viewer (2.5.0, Akoya Biosciences) software for annotation followed by capturing at high magnification. The Inform Viewer was used to visualize a detailed composite image of each section with the flexibility of spectral separation of all the fluorophores from each panel. StarDist (https://github.com/stardist/stardist), a deep learning tool extension available in the QuPath software (https://qupath.github.io, v0.40) was used to further analyze the expression of markers within a region of interest (ROI)^63^. Positive staining thresholds were manually defined for each marker following confirmation using the DAPI nuclear stain channel. Immune cell infiltration was evaluated in 10 random annotated ROI adjacent to the basal layer of the urothelium in the bladders for each treatment group. Composite object classifier tool was used to identify co-expression of markers. Measurements were calculated per mm^2^ and exported to GraphPad Prism (v9.5.1, California, USA) to analyze differences between sexes and treatment groups.

Single cell suspensions from spleens harvested were prepared by mechanical dissociation. Bone marrow was aspirated with RPMI-1640 media (Wisent Bioproducts, ON, Canada) using a 26-gauge needle and passing it through a 40 μm cell strainer followed by red blood cell (RBC) lysis. Details of further processing and multiparametric flow cytometry (Cytoflex-S; Beckman, USA) are provided in the **Supplementary methods** section.

#### Gene expression profiling of bladder using NanoString nCounter gene expression platform

Gene expression profiling was performed on total RNA isolated from fresh frozen whole bladders using a custom designed nCounter gene panel (**Suppl. Table S7)** which included genes associated with immune function, various cancer-related chemokines and cytokines, and housekeeping genes (NanoString Technologies, Washington, USA). Because of the significant changes observed in urothelium of female bladders following B cell depletion during BCG treatment, this analysis was only performed in bladders from female mice. Total RNA from fresh frozen bladders (as indicated) was isolated using the RNeasy Mini Kit (Cat. #74004, Qiagen, Hilden, Germany) as per the manufacturer’s instructions. The purity and concentration of isolated RNA was assessed using the NanoDrop ND-100 spectrophotometer (NanoDrop Technologies, DE, USA). Total RNA (150 ng) was used as a template for digital multiplexed profiling at Ontario Institute for Cancer Research as per previously established protocols. The nSolver software (NanoString Technologies, Washington, USA) was used to normalize the raw nCounter NanoString counts using built-in positive controls. Normalization was performed using housekeeping genes and overall assay efficiency was calculated using geometric mean of each control. Normalized nCounter gene counts between different treatment groups was further analysed in GraphPad Prism.

#### Plasma cytokine profiling and immunoglobulin isotyping

Female and male mice from each treatment group (n = 4-5 mice per group) were subjected to blood collection from the submandibular (facial) vein. Separated plasma was aliquoted as per the requirements for multiplex cytokine analysis and stored at -80°C until further profiling. Plasma was diluted using PBS (1:1) and subjected to multiplex cytokine analysis using the commercially available 32-Plex Mouse Cytokine Discovery Array (MD-31; Eve Technologies, AB, Canada).

Plasma Ig were analysed using the LEGENDplex Mouse Immunoglobulin Isotyping panel (6-plex) by flow cytometry (Cat #740493, BioLegend, CA, USA). Plasma sample and standards was prepared using assay buffer as per the manufacturer’s instruction. Data was analysed using the cloud-specific LEGENDplex Data Analysis Software Suite Qognit (v.2023.02.15, CA, USA). Statistically significant differences in the levels of Ig isotypes between the groups were determined using GraphPad Prism.

### *In vitro* studies

#### Cell culture

Splenic B cells were isolated from 12-month-old female and male mice (n=3-4/group) after 7 weeks of BBN exposure using EasySep Mouse Pan B cell negative isolation kit (Cat# 19844, StemCell technologies, BC, Canada). The purity and viability of enriched B cells was >90% as assessed by anti-B220 and anti-CD19 mAb staining using flow cytometry (**Fig. 5A; Fig. S5A).** B cells were resuspended in a base B cell culture media comprising of RPMI media supplemented with L-glutamine, 10% FBS, 1% Penicillin/Streptomycin, 1X Sodium Pyruvate, 50 µM 2-mercaptoethanol, 1X Nonessential amino acid and 10mM HEPES. 1.5 x 10^6^ cells/ml were seeded in a 96 well plate and treated with IFN-*γ* (20ng/ml, Cat#315-05, Peprotech, NJ, USA), IL-21 (20 ng/ml, Cat# 210-21, Peprotech, NJ, USA) and TLR7 agonist – Imiquimod (R837) (2.5µM, Cat# tlrl- imqs, Invivogen, CA, USA) for 6 and 18 hours at 37°C and 5% CO_2_. Cells were also infected with BCG TICE (Merck, Canada) along with IFN-*γ* and IL-21 to compare the expansion of ABCs. Lyophilized BCG was reconstituted in B cell culture media at a concentration of 1 mg/ml, resulting in approximately 0.2 – 1.6 x 10^7^ CFU per ml. Following an 18-hour treatment, the supernatant was collected, and cells were subjected to flow cytometric analysis using the ABC panel (**Supplementary methods**).

#### Quantitative real-time PCR

After 6 hrs of *in vitro* treatment, total RNA was extracted from the B cells using the RNeasy Mini Kit (Cat. #74004, QIAGEN, Germany) as per the manufacturer’s instruction. The quality and concentration of the RNA was assessed by the NanoDrop ND-100 spectrophotometer (NanoDrop Technologies, DE, USA). 150 ng of total RNA per sample was used to synthesize cDNA using LunaScript RT SuperMix kit (Cat# E3010L, NewEngland BioLabs, MA, USA) in a 20 µl reaction volume. Quantitative real-time PCR was performed using ViiA 7 Real-Time PCR System (Applied Biosystems, MA, USA). Briefly, the 20 µl reaction volume consisted of 10 µl of TaqMan Fast Advanced Master Mix (Cat# 4444556, Applied Biosystems, MA, USA), 2 µl of cDNA, 1 µl of TaqMan primers (**Suppl. Table S6)** and 7 µl of nuclease free water. Relative gene expression was calculated using 2^-ΔΔCt^ method and *Ubc* as the housekeeping gene.

#### Multiplex cytokine profiling of culture supernatants

Supernatants collected after 18 hours of treatment was aliquoted in separate tubes and stored at -80°C until further profiling. Supernatant was subjected to multiplex cytokine analysis using commercially available Mouse Cytokine Proinflammatory Focused 10-plex Discovery Array (MD-F10; Eve Technologies, AB, Canada).

#### Immunocytochemistry

Splenic B cells, enriched *via* negative selection, were treated with IFN-γ, IL-21, TLR7, and BCG infection for 6 hours and 18 hours. The treated cells were then transferred to cover slips pre-coated with a 0.1% Poly-L-lysine solution (Cat# P8920, Millipore Sigma, MO, USA) and incubated at 37°C with 5% CO_2_ for 1 hour. Next, the cells were fixed with 4% paraformaldehyde (PFA) for 30 minutes at room temperature (RT) and blocked with 5% skim milk for 30–45 minutes at RT. Subsequently, the cells were incubated with primary antibodies Anti-CD11c and Anti-LAM (**Suppl. Table S5**) overnight at 4°C. The cells were then incubated with the corresponding secondary antibodies (**Suppl. Table S5**) for 1 hour at RT. All intermediate washes were performed using PBS-T (0.1% Tween 20) buffer. Finally, ProLong Gold Antifade Mountant with DNA stain DAPI (Cat# P36941, Invitrogen, MA, USA) was used to prevent photobleaching, and the cells were transferred onto slides. The slides were visualized using a Zeiss Axio Imager M2 epifluorescence microscope (Zeiss, Germany).

### Statistical Analysis

Statistical analysis was performed using GraphPad Prism. Results are expressed as mean ± SD. Comparisons between two groups with one independent variable was performed using non-parametric t-test (Mann-Whitney). Three or more groups with one independent variable were analysed using one-way ANOVA with Tukey’s post hoc test. Three or more groups with two or more independent variable were analysed using two-way ANOVA with Tukey’s multiple comparison test. Statistical significance is indicated by *p<0.05, **p<0.01, ***p<0.001, ****p<0.001, ns - not significant.

## Author contributions and acknowledgements

MK conceptualized and designed the study. PY, SR, SC, GC and MK performed experiments included in this study, analyzed data and contributed to manuscript writing and reviewing. IE helped with BBN experiments. KS and JH analyzed the NanoString GeoMx DSP data. DJ helped with chart review and identification of BCG responders and non-responders. MX helped with reviewing histopathological features and retrieving the archival tumor tissue specimens. DRS and DMB helped with clinical classifications of BCG treated patient specimens. EM and JM designed the Nanostring nCounter panel in consultation with WK. All authors reviewed the manuscript. This study was supported by research operating grants from Bladder Cancer Canada, Cancer Research Society, Ontario Ministry of Research Innovation and Science: Early Researcher Award and Canada Foundation for Innovation, to MK. We thank Lee Boudreau at QLMP for his assistance with embedding of FFPE blocks, sectioning, and H&E staining. We thank Shakeel Virk at the QLMP for his assistance with imaging and Halo software and Katy Milne at BC Cancer’s Molecular and Cellular Immunology Core (MCIC) for immunofluorescence staining of TMAs. Dr. Patricia Lima at the Queen’s Cardiopulmonary Unit helped with optimization of ABC panel and confocal microscopy-based imaging of multiplex IF stained whole bladder sections. We thank Dr. Andrew Winterborn and the staff at the Queen’s animal care facility for all the support with animal studies.

## SUPPLEMENTARY TABLES

**Supplementary Table S1.**
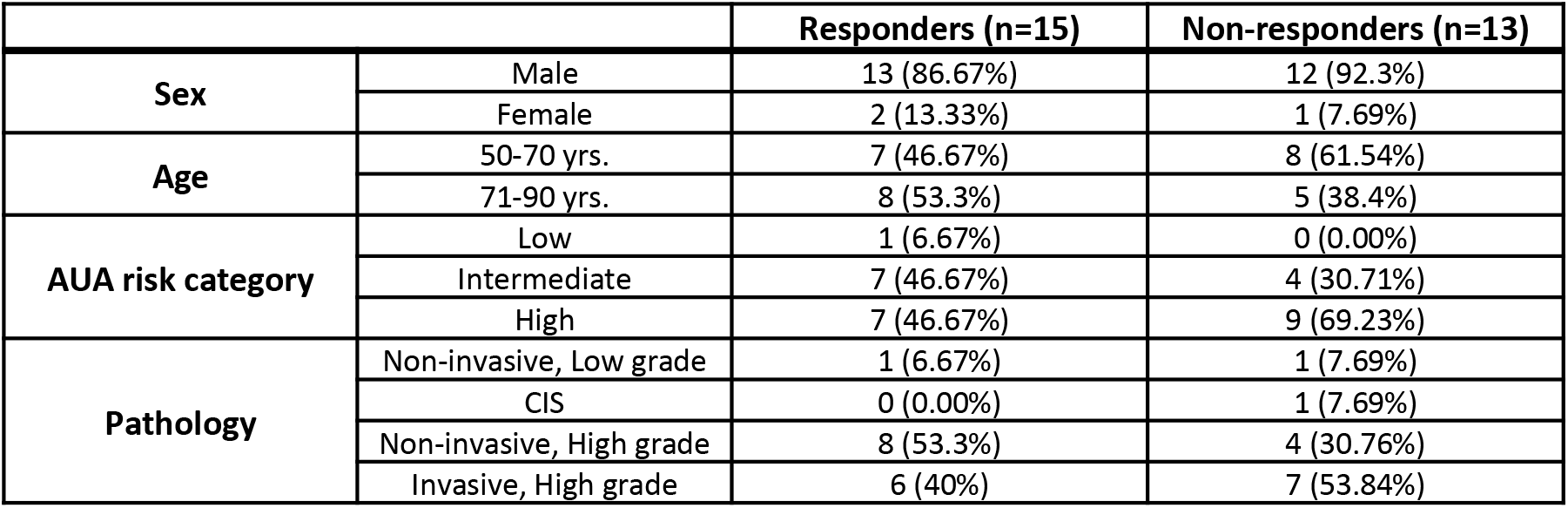
Patient clinical and pathological features.

**Supplementary Table S2.**
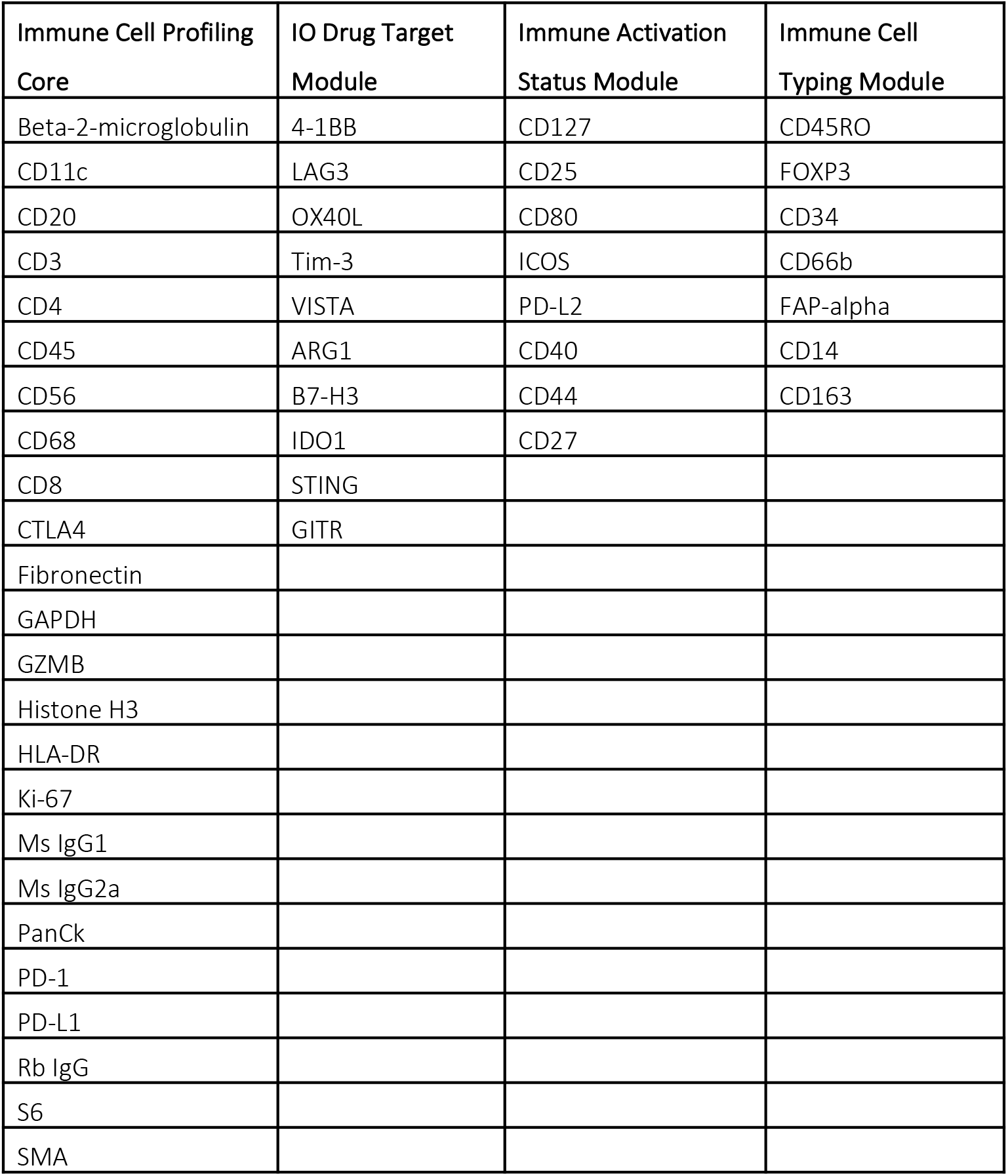
List of proteins investigated via NanoString GeoMx DSP.

**Supplementary Table S3.**
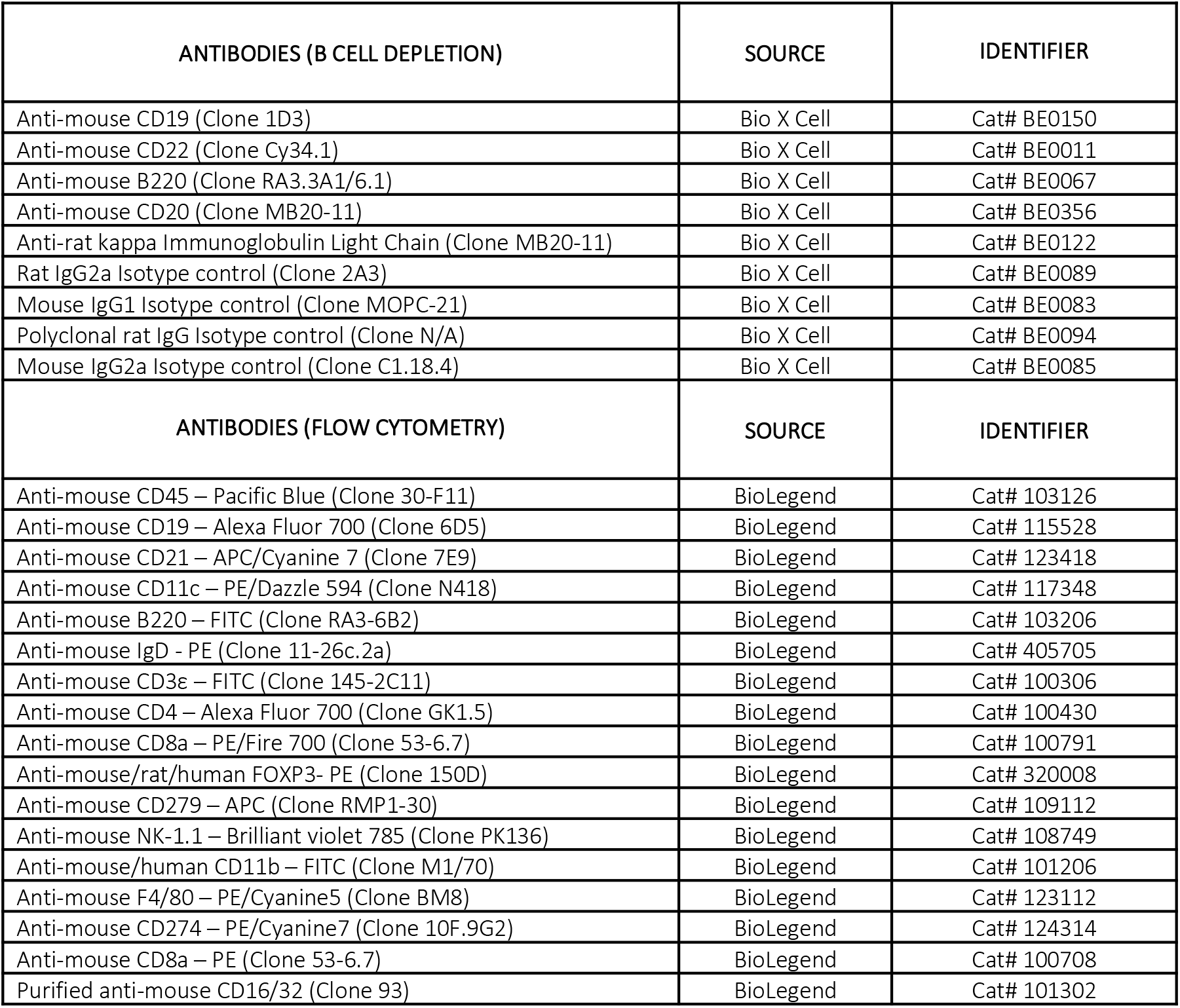
List of antibodies used in multispectral flow cytometry.

**Supplementary Table S4.**
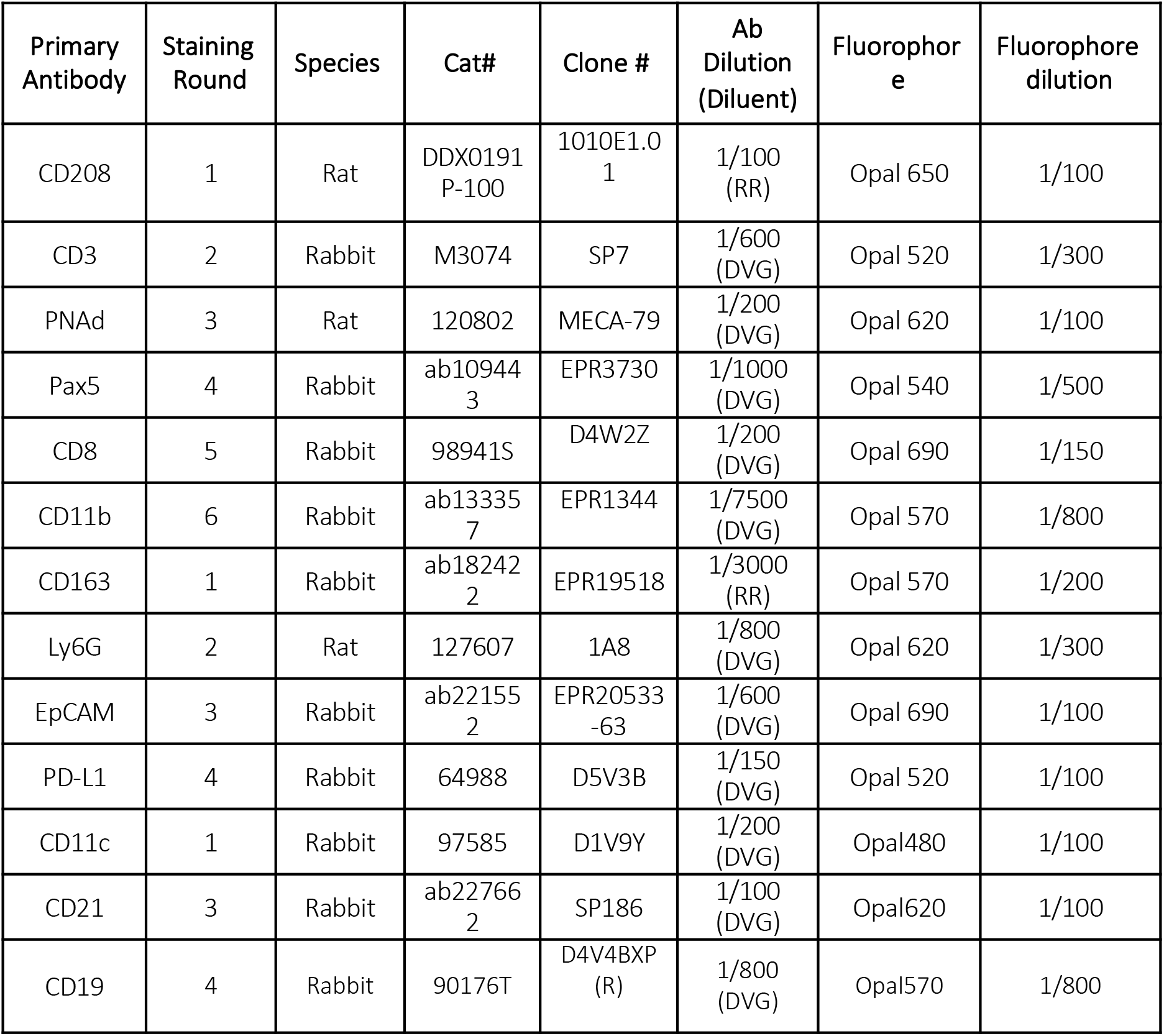
Antibodies used in multiplex IF staining.

**Supplementary Table S5.**
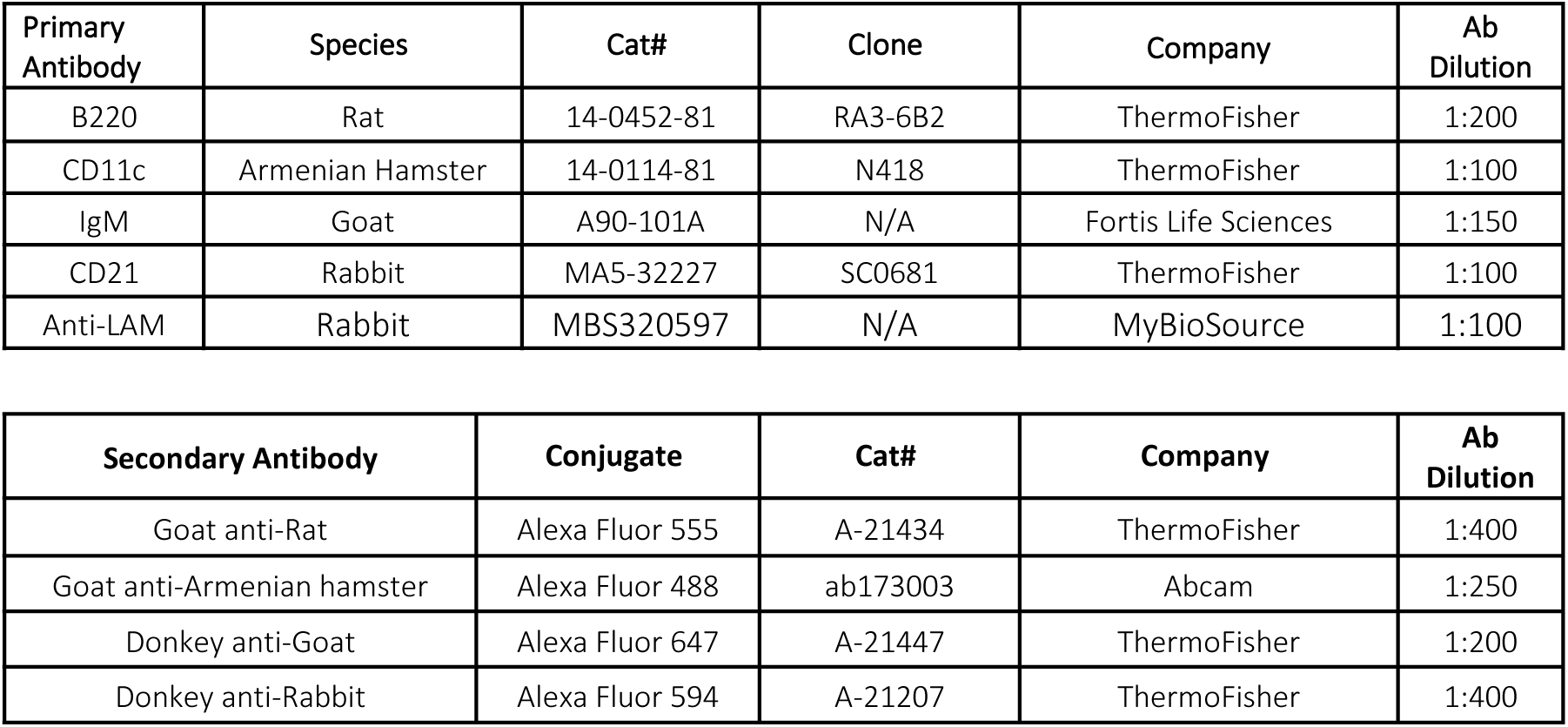
Antibodies used in atypical B cell panel for multiplex IF staining.

**Supplementary Table S6.**
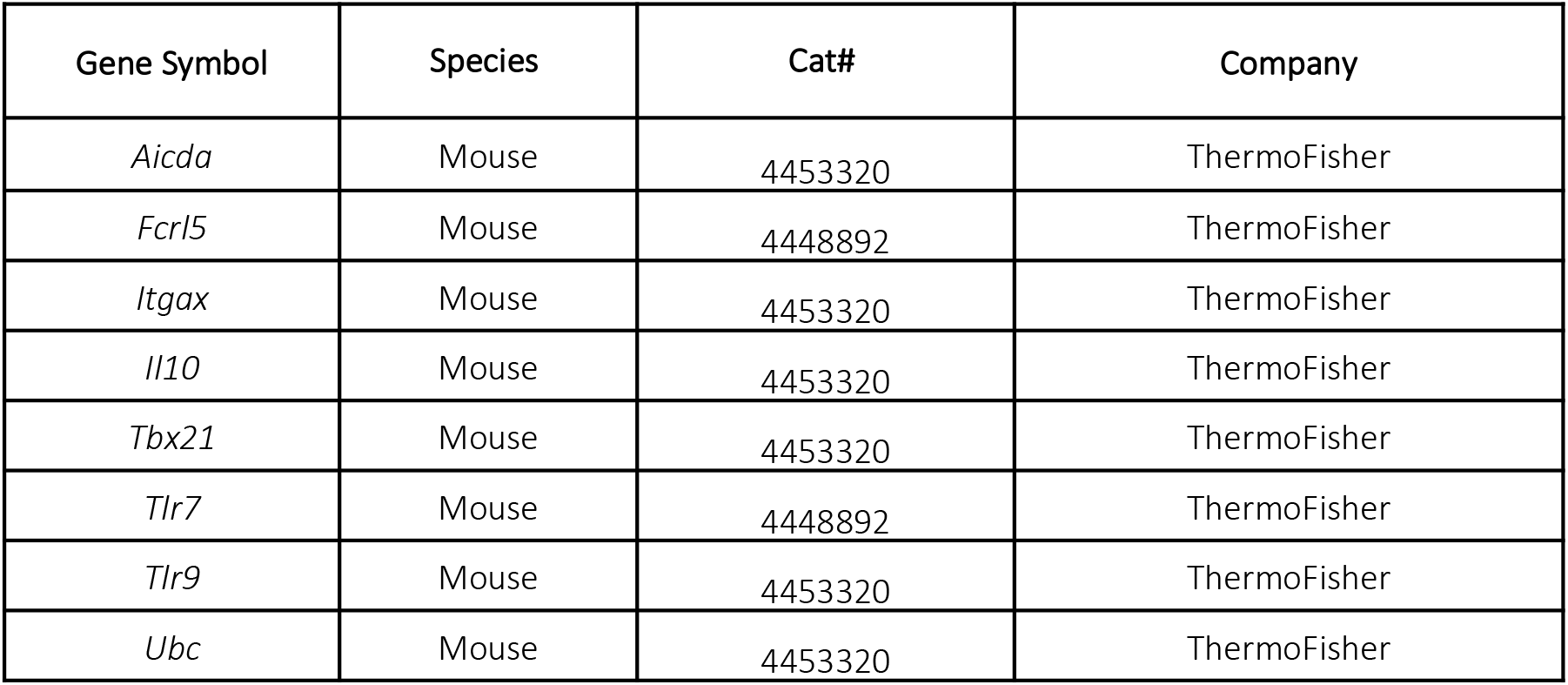
List of primers used for quantitative RT-PCR.

**Supplementary Table S7.**
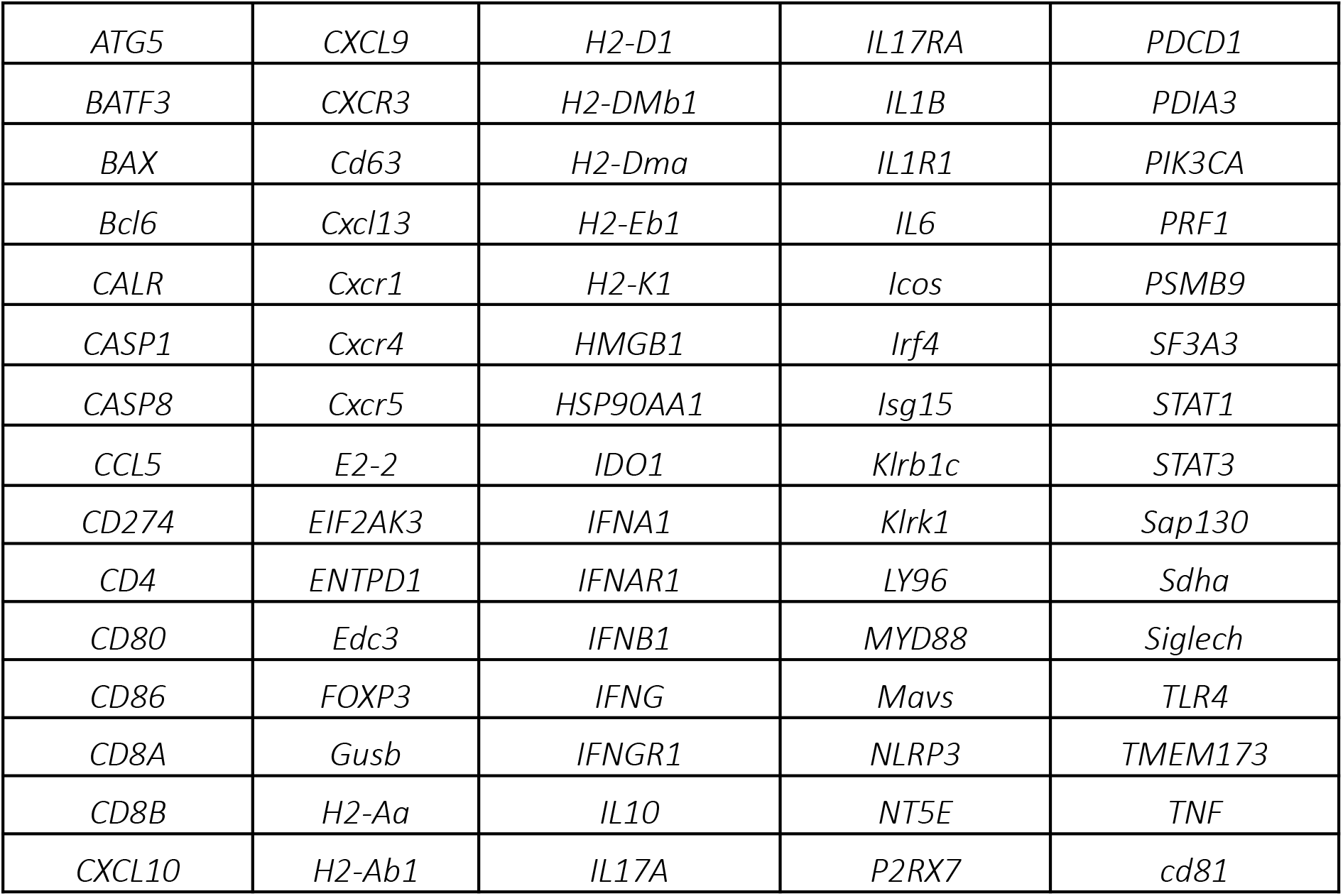
List of genes investigated via NanoString nCounter.

**Supplementary Table S8.**
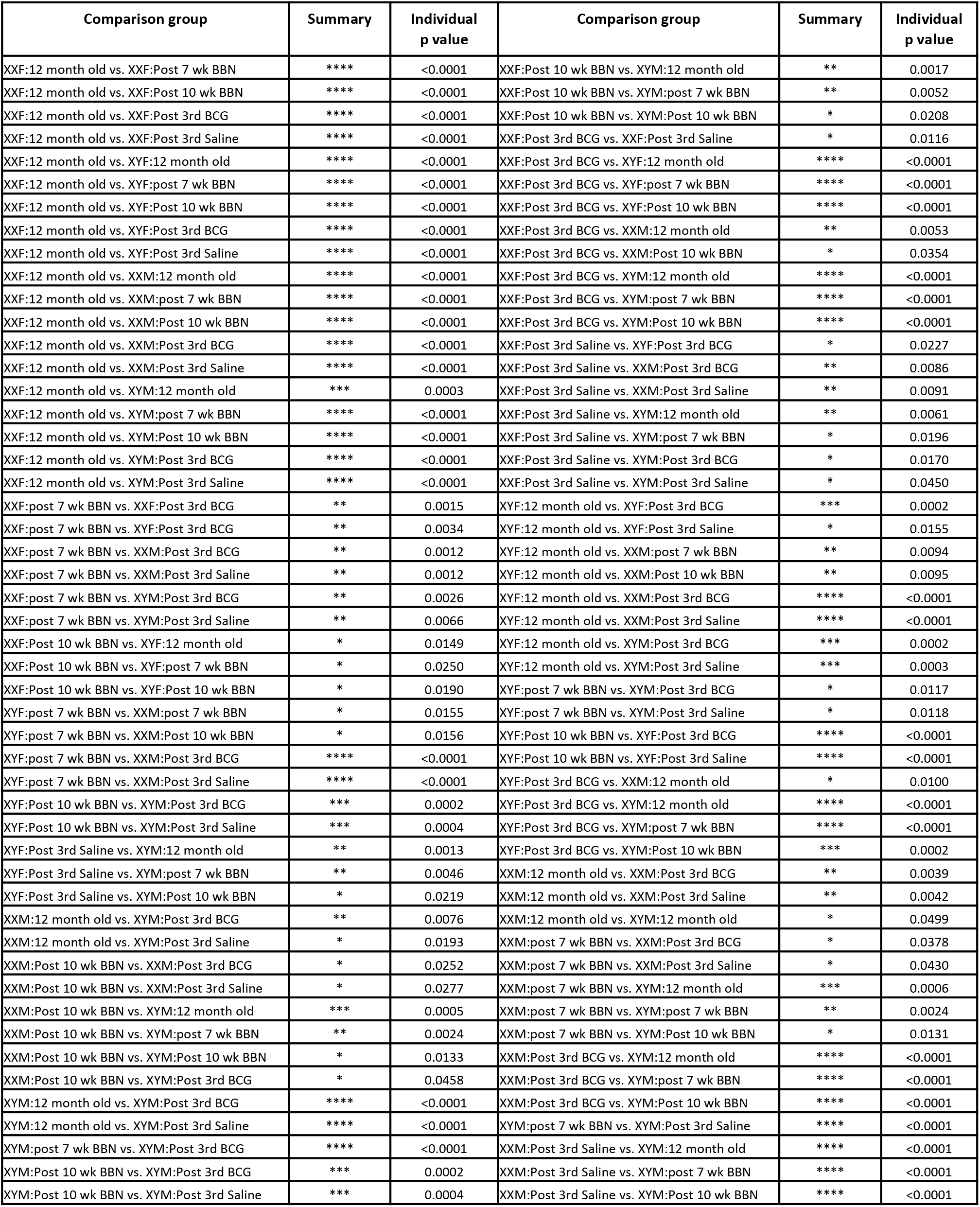
Summary of statistical comparison of splenic ABC profiles between the treatment groups of the four genotypes of FCG.

