## Supplementary figures for "Atypical B cells mediate poor response to Bacillus Calmette Guérin immunotherapy in non-muscle invasive bladder cancer"

### SUPPLEMENTARY TABLES

#### Supplementary Table S1. Patient clinical and pathological features

| | | Responders (n=15) | Non-responders (n=13) |
|---|---|---|---|
| **Sex** | Male | 13 (86.67%) | 12 (92.3%) |
| | Female | 2 (13.33%) | 1 (7.69%) |
| **Age** | 50-70 yrs. | 7 (46.67%) | 8 (61.54%) |
| | 71-90 yrs. | 8 (53.3%) | 5 (38.4%) |
| **AUA risk category** | Low | 1 (6.67%) | 0 (0.00%) |
| | Intermediate | 7 (46.67%) | 4 (30.71%) |
| | High | 7 (46.67%) | 9 (69.23%) |
| **Pathology** | Non-invasive, Low grade | 1 (6.67%) | 1 (7.69%) |
| | CIS | 0 (0.00%) | 1 (7.69%) |
| | Non-invasive, High grade | 8 (53.3%) | 4 (30.76%) |
| | Invasive, High grade | 6 (40%) | 7 (53.84%) |

#### Supplementary Table S2. List of proteins investigated via NanoString GeoMx DSP

| Immune Cell Profiling Core | IO Drug Target Module | Immune Activation Status Module | Immune Cell Typing Module |
|---|---|---|---|
| Beta-2-microglobulin | 4-1BB | CD127 | CD45RO |
| CD11c | LAG3 | CD25 | FOXP3 |
| CD20 | OX40L | CD80 | CD34 |
| CD3 | Tim-3 | ICOS | CD66b |
| CD4 | VISTA | PD-L2 | FAP-alpha |
| CD45 | ARG1 | CD40 | CD14 |
| CD56 | B7-H3 | CD44 | CD163 |
| CD68 | IDO1 | CD27 | |
| CD8 | STING | | |
| CTLA4 | GITR | | |
| Fibronectin | | | |
| GAPDH | | | |
| GZMB | | | |
| Histone H3 | | | |
| HLA-DR | | | |
| Ki-67 | | | |
| Ms IgG1 | | | |
| Ms IgG2a | | | |
| PanCk | | | |
| PD-1 | | | |
| PD-L1 | | | |
| Rb IgG | | | |
| S6 | | | |
| SMA | | | |

Supplementary Table S3. List of antibodies used in multispectral flow cytometry

| ANTIBODIES (B CELL DEPLETION) | SOURCE | IDENTIFIER |
|---|---|---|
| Anti-mouse CD19 (Clone 1D3) | Bio X Cell | Cat# BE0150 |
| Anti-mouse CD22 (Clone Cy34.1) | Bio X Cell | Cat# BE0011 |
| Anti-mouse B220 (Clone RA3.3A1/6.1) | Bio X Cell | Cat# BE0067 |
| Anti-mouse CD20 (Clone MB20-11) | Bio X Cell | Cat# BE0356 |
| Anti-rat kappa Immunoglobulin Light Chain (Clone MB20-11) | Bio X Cell | Cat# BE0122 |
| Rat IgG2a Isotype control (Clone 2A3) | Bio X Cell | Cat# BE0089 |
| Mouse IgG1 Isotype control (Clone MOPC-21) | Bio X Cell | Cat# BE0083 |
| Polyclonal rat IgG Isotype control (Clone N/A) | Bio X Cell | Cat# BE0094 |
| Mouse IgG2a Isotype control (Clone C1.18.4) | Bio X Cell | Cat# BE0085 |
| ANTIBODIES (FLOW CYTOMETRY) | SOURCE | IDENTIFIER |
| Anti-mouse CD45 – Pacific Blue (Clone 30-F11) | BioLegend | Cat# 103126 |
| Anti-mouse CD19 – Alexa Fluor 700 (Clone 6D5) | BioLegend | Cat# 115528 |
| Anti-mouse CD21 – APC/Cyanine 7 (Clone 7E9) | BioLegend | Cat# 123418 |
| Anti-mouse CD11c – PE/Dazzle 594 (Clone N418) | BioLegend | Cat# 117348 |
| Anti-mouse B220 – FITC (Clone RA3-6B2) | BioLegend | Cat# 103206 |
| Anti-mouse IgD - PE (Clone 11-26c.2a) | BioLegend | Cat# 405705 |
| Anti-mouse CD3ε – FITC (Clone 145-2C11) | BioLegend | Cat# 100306 |
| Anti-mouse CD4 – Alexa Fluor 700 (Clone GK1.5) | BioLegend | Cat# 100430 |
| Anti-mouse CD8a – PE/Fire 700 (Clone 53-6.7) | BioLegend | Cat# 100791 |
| Anti-mouse/rat/human FOXP3- PE (Clone 150D) | BioLegend | Cat# 320008 |
| Anti-mouse CD279 – APC (Clone RMP1-30) | BioLegend | Cat# 109112 |
| Anti-mouse NK-1.1 – Brilliant violet 785 (Clone PK136) | BioLegend | Cat# 108749 |
| Anti-mouse/human CD11b – FITC (Clone M1/70) | BioLegend | Cat# 101206 |
| Anti-mouse F4/80 – PE/Cyanine5 (Clone BM8) | BioLegend | Cat# 123112 |
| Anti-mouse CD274 – PE/Cyanine7 (Clone 10F.9G2) | BioLegend | Cat# 124314 |
| Anti-mouse CD8a – PE (Clone 53-6.7) | BioLegend | Cat# 100708 |
| Purified anti-mouse CD16/32 (Clone 93) | BioLegend | Cat# 101302 |

Supplementary Table S4. Antibodies used in multiplex IF staining.

| Primary Antibody | Staining Round | Species | Cat# | Clone # | Ab Dilution (Diluent) | Fluorophore | Fluorophore dilution |
|---|---|---|---|---|---|---|---|
| CD208 | 1 | Rat | DDX0191P-100 | 1010E1.01 | 1/100 (RR) | Opal 650 | 1/100 |
| CD3 | 2 | Rabbit | M3074 | SP7 | 1/600 (DVG) | Opal 520 | 1/300 |
| PNAd | 3 | Rat | 120802 | MECA-79 | 1/200 (DVG) | Opal 620 | 1/100 |
| Pax5 | 4 | Rabbit | ab109443 | EPR3730 | 1/1000 (DVG) | Opal 540 | 1/500 |
| CD8 | 5 | Rabbit | 98941S | D4W2Z | 1/200 (DVG) | Opal 690 | 1/150 |
| CD11b | 6 | Rabbit | ab133357 | EPR1344 | 1/7500 (DVG) | Opal 570 | 1/800 |
| CD163 | 1 | Rabbit | ab182422 | EPR19518 | 1/3000 (RR) | Opal 570 | 1/200 |
| Ly6G | 2 | Rat | 127607 | 1A8 | 1/800 (DVG) | Opal 620 | 1/300 |
| EpCAM | 3 | Rabbit | ab221552 | EPR20533-63 | 1/600 (DVG) | Opal 690 | 1/100 |
| PD-L1 | 4 | Rabbit | 64988 | D5V3B | 1/150 (DVG) | Opal 520 | 1/100 |
| CD11c | 1 | Rabbit | 97585 | D1V9Y | 1/200 (DVG) | Opal480 | 1/100 |
| CD21 | 3 | Rabbit | ab227662 | SP186 | 1/100 (DVG) | Opal620 | 1/100 |
| CD19 | 4 | Rabbit | 90176T | D4V4BXP (R) | 1/800 (DVG) | Opal570 | 1/800 |

Supplementary Table S5. Antibodies used in atypical B cell panel for multiplex IF staining.

| Primary Antibody | Species | Cat# | Clone | Company | Ab Dilution |
|---|---|---|---|---|---|
| B220 | Rat | 14-0452-81 | RA3-6B2 | ThermoFisher | 1:200 |
| CD11c | Armenian Hamster | 14-0114-81 | N418 | ThermoFisher | 1:100 |
| IgM | Goat | A90-101A | N/A | Fortis Life Sciences | 1:150 |
| CD21 | Rabbit | MA5-32227 | SC0681 | ThermoFisher | 1:100 |
| Anti-LAM | Rabbit | MBS320597 | N/A | MyBioSource | 1:100 |

| **Secondary Antibody** | **Conjugate** | **Cat#** | **Company** | **Ab Dilution** |
|---|---|---|---|---|
| Goat anti-Rat | Alexa Fluor 555 | A-21434 | ThermoFisher | 1:400 |
| Goat anti-Armenian hamster | Alexa Fluor 488 | ab173003 | Abcam | 1:250 |
| Donkey anti-Goat | Alexa Fluor 647 | A-21447 | ThermoFisher | 1:200 |
| Donkey anti-Rabbit | Alexa Fluor 594 | A-21207 | ThermoFisher | 1:400 |

Supplementary Table S6. List of primers used for quantitative RT-PCR

| Gene Symbol | Species | Cat# | Company |
|---|---|---|---|
| *Aicda* | Mouse | 4453320 | ThermoFisher |
| *Fcrl5* | Mouse | 4448892 | ThermoFisher |
| *Itgax* | Mouse | 4453320 | ThermoFisher |
| *Il10* | Mouse | 4453320 | ThermoFisher |
| *Tbx21* | Mouse | 4453320 | ThermoFisher |
| *Tlr7* | Mouse | 4448892 | ThermoFisher |
| *Tlr9* | Mouse | 4453320 | ThermoFisher |
| *Ubc* | Mouse | 4453320 | ThermoFisher |

Supplementary Table S7. List of genes investigated via NanoString nCounter

| | | | | |
|---|---|---|---|---|
| *ATG5* | *CXCL9* | *H2-D1* | *IL17RA* | *PDCD1* |
| *BATF3* | *CXCR3* | *H2-DMb1* | *IL1B* | *PDIA3* |
| *BAX* | *Cd63* | *H2-Dma* | *IL1R1* | *PIK3CA* |
| *Bcl6* | *Cxcl13* | *H2-Eb1* | *IL6* | *PRF1* |
| *CALR* | *Cxcr1* | *H2-K1* | *Icos* | *PSMB9* |
| *CASP1* | *Cxcr4* | *HMGB1* | *Irf4* | *SF3A3* |
| *CASP8* | *Cxcr5* | *HSP90AA1* | *Isg15* | *STAT1* |
| *CCL5* | *E2-2* | *IDO1* | *Klrb1c* | *STAT3* |
| *CD274* | *EIF2AK3* | *IFNA1* | *Klrk1* | *Sap130* |
| *CD4* | *ENTPD1* | *IFNAR1* | *LY96* | *Sdha* |
| *CD80* | *Edc3* | *IFNB1* | *MYD88* | *Siglech* |
| *CD86* | *FOXP3* | *IFNG* | *Mavs* | *TLR4* |
| *CD8A* | *Gusb* | *IFNGR1* | *NLRP3* | *TMEM173* |
| *CD8B* | *H2-Aa* | *IL10* | *NT5E* | *TNF* |
| *CXCL10* | *H2-Ab1* | *IL17A* | *P2RX7* | *cd81* |

Supplementary Table S8. Summary of statistical comparison of splenic ABC profiles between the treatment groups of the four genotypes of FCG

| Comparison group | Summary | Individual p value | Comparison group | Summary | Individual p value |
|---|---|---|---|---|---|
| XXF:12 month old vs. XXF:Post 7 wk BBN | **** | <0.0001 | XXF:Post 10 wk BBN vs. XYM:12 month old | ** | 0.0017 |
| XXF:12 month old vs. XXF:Post 10 wk BBN | **** | <0.0001 | XXF:Post 10 wk BBN vs. XYM:post 7 wk BBN | ** | 0.0052 |
| XXF:12 month old vs. XXF:Post 3rd BCG | **** | <0.0001 | XXF:Post 10 wk BBN vs. XYM:Post 10 wk BBN | * | 0.0208 |
| XXF:12 month old vs. XXF:Post 3rd Saline | **** | <0.0001 | XXF:Post 3rd BCG vs. XXF:Post 3rd Saline | * | 0.0116 |
| XXF:12 month old vs. XYF:12 month old | **** | <0.0001 | XXF:Post 3rd BCG vs. XYF:12 month old | **** | <0.0001 |
| XXF:12 month old vs. XYF:post 7 wk BBN | **** | <0.0001 | XXF:Post 3rd BCG vs. XYF:post 7 wk BBN | **** | <0.0001 |
| XXF:12 month old vs. XYF:Post 10 wk BBN | **** | <0.0001 | XXF:Post 3rd BCG vs. XYF:Post 10 wk BBN | **** | <0.0001 |
| XXF:12 month old vs. XYF:Post 3rd BCG | **** | <0.0001 | XXF:Post 3rd BCG vs. XXM:12 month old | ** | 0.0053 |
| XXF:12 month old vs. XYF:Post 3rd Saline | **** | <0.0001 | XXF:Post 3rd BCG vs. XXM:Post 10 wk BBN | * | 0.0354 |
| XXF:12 month old vs. XXM:12 month old | **** | <0.0001 | XXF:Post 3rd BCG vs. XYM:12 month old | **** | <0.0001 |
| XXF:12 month old vs. XXM:post 7 wk BBN | **** | <0.0001 | XXF:Post 3rd BCG vs. XYM:post 7 wk BBN | **** | <0.0001 |
| XXF:12 month old vs. XXM:Post 10 wk BBN | **** | <0.0001 | XXF:Post 3rd BCG vs. XYM:Post 10 wk BBN | **** | <0.0001 |
| XXF:12 month old vs. XXM:Post 3rd BCG | **** | <0.0001 | XXF:Post 3rd Saline vs. XYF:Post 3rd BCG | * | 0.0227 |
| XXF:12 month old vs. XXM:Post 3rd Saline | **** | <0.0001 | XXF:Post 3rd Saline vs. XXM:Post 3rd BCG | ** | 0.0086 |
| XXF:12 month old vs. XYM:12 month old | *** | 0.0003 | XXF:Post 3rd Saline vs. XXM:Post 3rd Saline | ** | 0.0091 |
| XXF:12 month old vs. XYM:post 7 wk BBN | **** | <0.0001 | XXF:Post 3rd Saline vs. XYM:12 month old | ** | 0.0061 |
| XXF:12 month old vs. XYM:Post 10 wk BBN | **** | <0.0001 | XXF:Post 3rd Saline vs. XYM:post 7 wk BBN | * | 0.0196 |
| XXF:12 month old vs. XYM:Post 3rd BCG | **** | <0.0001 | XXF:Post 3rd Saline vs. XYM:Post 3rd BCG | * | 0.0170 |
| XXF:12 month old vs. XYM:Post 3rd Saline | **** | <0.0001 | XXF:Post 3rd Saline vs. XYM:Post 3rd Saline | * | 0.0450 |
| XXF:post 7 wk BBN vs. XXF:Post 3rd BCG | ** | 0.0015 | XYF:12 month old vs. XYF:Post 3rd BCG | *** | 0.0002 |
| XXF:post 7 wk BBN vs. XYF:Post 3rd BCG | ** | 0.0034 | XYF:12 month old vs. XYF:Post 3rd Saline | * | 0.0155 |
| XXF:post 7 wk BBN vs. XXM:Post 3rd BCG | ** | 0.0012 | XYF:12 month old vs. XXM:post 7 wk BBN | ** | 0.0094 |
| XXF:post 7 wk BBN vs. XXM:Post 3rd Saline | ** | 0.0012 | XYF:12 month old vs. XXM:Post 10 wk BBN | ** | 0.0095 |
| XXF:post 7 wk BBN vs. XYM:Post 3rd BCG | ** | 0.0026 | XYF:12 month old vs. XXM:Post 3rd BCG | **** | <0.0001 |
| XXF:post 7 wk BBN vs. XYM:Post 3rd Saline | ** | 0.0066 | XYF:12 month old vs. XXM:Post 3rd Saline | **** | <0.0001 |
| XXF:Post 10 wk BBN vs. XYF:12 month old | * | 0.0149 | XYF:12 month old vs. XYM:Post 3rd BCG | *** | 0.0002 |
| XXF:Post 10 wk BBN vs. XYF:post 7 wk BBN | * | 0.0250 | XYF:12 month old vs. XYM:Post 3rd Saline | *** | 0.0003 |
| XXF:Post 10 wk BBN vs. XYF:Post 10 wk BBN | * | 0.0190 | XYF:post 7 wk BBN vs. XYM:Post 3rd BCG | * | 0.0117 |
| XYF:post 7 wk BBN vs. XXM:post 7 wk BBN | * | 0.0155 | XYF:post 7 wk BBN vs. XYM:Post 3rd Saline | * | 0.0118 |
| XYF:post 7 wk BBN vs. XXM:Post 10 wk BBN | * | 0.0156 | XYF:Post 10 wk BBN vs. XYF:Post 3rd BCG | **** | <0.0001 |
| XYF:post 7 wk BBN vs. XXM:Post 3rd BCG | **** | <0.0001 | XYF:Post 10 wk BBN vs. XYF:Post 3rd Saline | **** | <0.0001 |
| XYF:post 7 wk BBN vs. XXM:Post 3rd Saline | **** | <0.0001 | XYF:Post 3rd BCG vs. XXM:12 month old | * | 0.0100 |
| XYF:Post 10 wk BBN vs. XYM:Post 3rd BCG | *** | 0.0002 | XYF:Post 3rd BCG vs. XYM:12 month old | **** | <0.0001 |
| XYF:Post 10 wk BBN vs. XYM:Post 3rd Saline | *** | 0.0004 | XYF:Post 3rd BCG vs. XYM:post 7 wk BBN | **** | <0.0001 |
| XYF:Post 3rd Saline vs. XYM:12 month old | ** | 0.0013 | XYF:Post 3rd BCG vs. XYM:Post 10 wk BBN | *** | 0.0002 |
| XYF:Post 3rd Saline vs. XYM:post 7 wk BBN | ** | 0.0046 | XXM:12 month old vs. XXM:Post 3rd BCG | ** | 0.0039 |
| XYF:Post 3rd Saline vs. XYM:Post 10 wk BBN | * | 0.0219 | XXM:12 month old vs. XXM:Post 3rd Saline | ** | 0.0042 |
| XXM:12 month old vs. XYM:Post 3rd BCG | ** | 0.0076 | XXM:12 month old vs. XYM:12 month old | * | 0.0499 |
| XXM:12 month old vs. XYM:Post 3rd Saline | * | 0.0193 | XXM:post 7 wk BBN vs. XXM:Post 3rd BCG | * | 0.0378 |
| XXM:Post 10 wk BBN vs. XXM:Post 3rd BCG | * | 0.0252 | XXM:post 7 wk BBN vs. XXM:Post 3rd Saline | * | 0.0430 |
| XXM:Post 10 wk BBN vs. XXM:Post 3rd Saline | * | 0.0277 | XXM:post 7 wk BBN vs. XYM:12 month old | *** | 0.0006 |
| XXM:Post 10 wk BBN vs. XYM:12 month old | *** | 0.0005 | XXM:post 7 wk BBN vs. XYM:post 7 wk BBN | ** | 0.0024 |
| XXM:Post 10 wk BBN vs. XYM:post 7 wk BBN | ** | 0.0024 | XXM:post 7 wk BBN vs. XYM:Post 10 wk BBN | * | 0.0131 |
| XXM:Post 10 wk BBN vs. XYM:Post 10 wk BBN | * | 0.0133 | XXM:Post 3rd BCG vs. XYM:12 month old | **** | <0.0001 |
| XXM:Post 10 wk BBN vs. XYM:Post 3rd BCG | * | 0.0458 | XXM:Post 3rd BCG vs. XYM:post 7 wk BBN | **** | <0.0001 |
| XYM:12 month old vs. XYM:Post 3rd BCG | **** | <0.0001 | XXM:Post 3rd BCG vs. XYM:Post 10 wk BBN | **** | <0.0001 |
| XYM:12 month old vs. XYM:Post 3rd Saline | **** | <0.0001 | XYM:post 7 wk BBN vs. XYM:Post 3rd Saline | **** | <0.0001 |
| XYM:post 7 wk BBN vs. XYM:Post 3rd BCG | **** | <0.0001 | XXM:Post 3rd Saline vs. XYM:12 month old | **** | <0.0001 |
| XYM:Post 10 wk BBN vs. XYM:Post 3rd BCG | *** | 0.0002 | XXM:Post 3rd Saline vs. XYM:post 7 wk BBN | **** | <0.0001 |
| XYM:Post 10 wk BBN vs. XYM:Post 3rd Saline | *** | 0.0004 | XXM:Post 3rd Saline vs. XYM:Post 10 wk BBN | **** | <0.0001 |

### SUPPLEMENTARY FIGURES

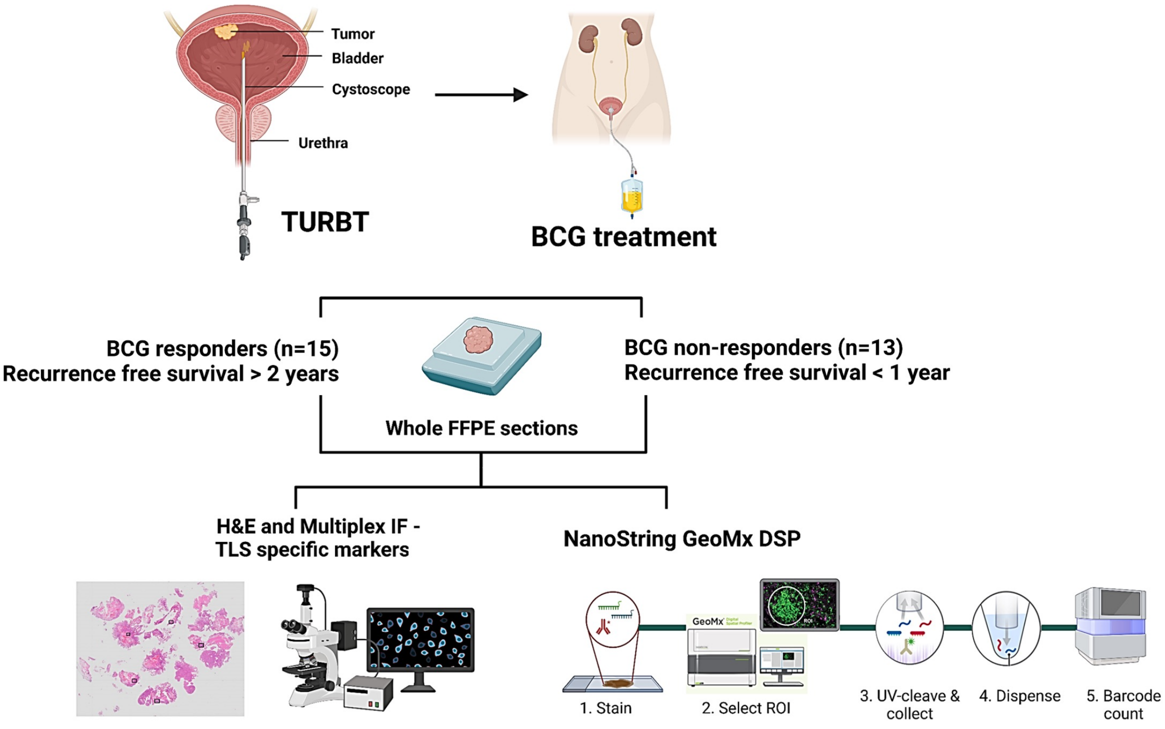


**Figure S1.** Schematic showing workflow for characterization of TLSs using multiplex Immunofluorescence and NanoString GeoMx DSP based profiling

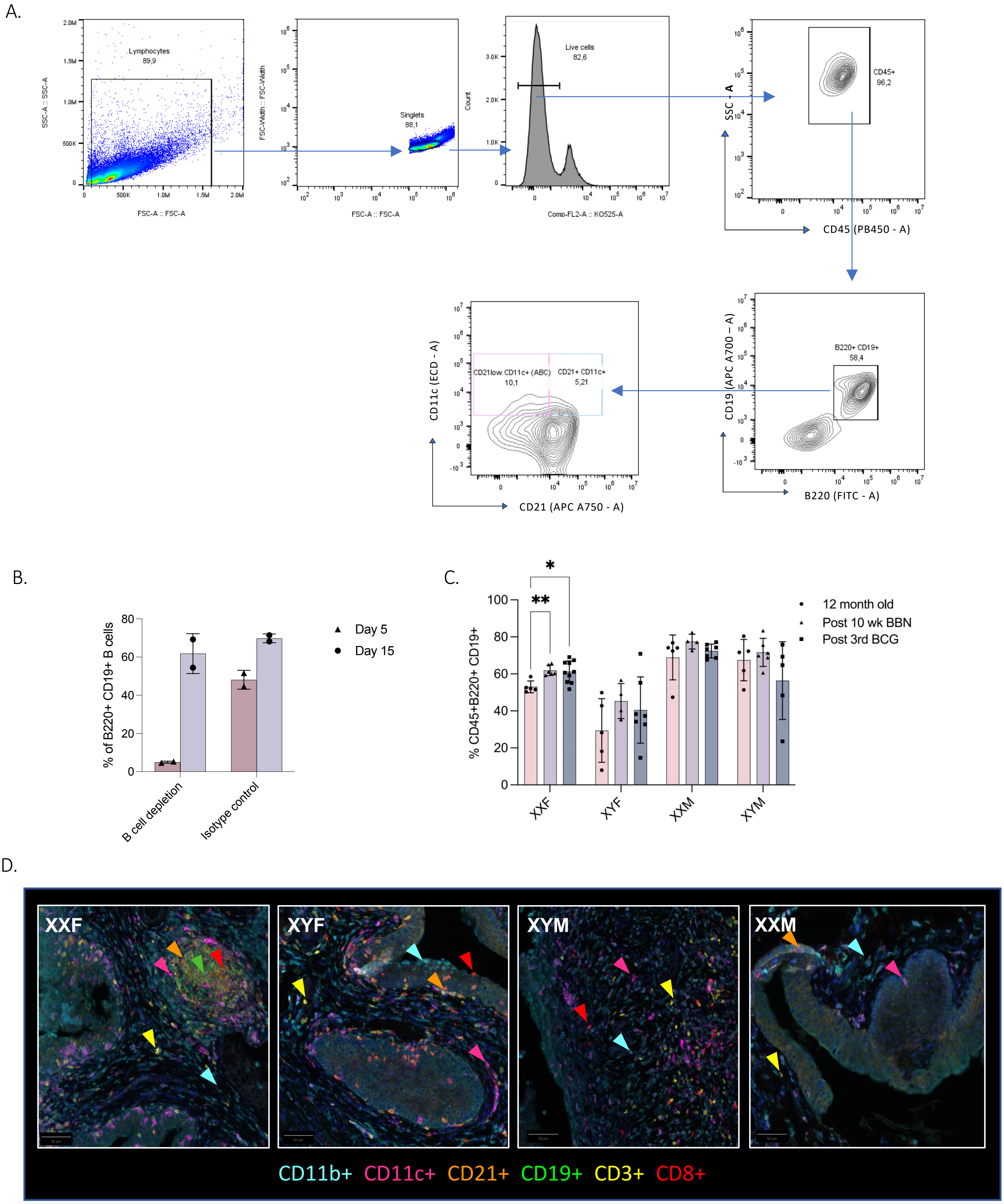

A.
Lymphocytes
89,9
SSC-A :: SSC-A
FSC-A :: FSC-A
Singlets
88,1
FSC-Width :: FSC-Width
Live cells
82,6
Count
Comp-FL2-A :: KO525-A
CD45+
96,2
SSC - A
CD45 (PB450 - A)
B220+ CD19+
58,4
CD19 (APC A700 – A)
B220 (FITC - A)
CD21low CD11c+ (ABC)
10,1
CD21+ CD11c+
5,21
CD11c (ECD - A)
CD21 (APC A750 - A)
B.
% of B220+ CD19+ B cells
Day 5
Day 15
B cell depletion
Isotype control
C.
% CD45+B220+ CD19+
12 month old
Post 10 wk BBN
Post 3rd BCG
XXF
XYF
XXM
XYM
D.
XXF
XYF
XYM
XXM
CD11b+ CD11c+ CD21+ CD19+ CD3+ CD8+

**Figure S2. Gating strategy for identifying atypical B cells (ABCs).** Single cell suspensions from spleen and bone marrow of mice were stained with ABC panel antibodies and subjected to multispectral flow cytometric analysis. Single cells was first identified by plotting forward scatter width (FSC-Width) against side scatter area (SSC-A). Events deviating from the linear cell population, called as doublets were excluded. Live cells were evaluated by gating on viability dye-negative cell. CD45 was used to discriminate the total immune cell population. B220+ CD19+ cells revealed the proportion of total B cells. Further plotting against CD21 and CD11c markers was used to identify ABC population (identified as CD21-/low CD11c+) (A). Single color (SC) and fluorescence minus one (FMO) was used to determine the position of gating for all subsets and further compared with unstained samples. All analysis were done in FlowJo® software. Bar graph represents number of total B220+ CD19+ total B cells in 9-month-old female mice in the depleted and isotype control group at day 5 and day 15 post treatment (n=2, B). Bar graph showing splenic B220+ CD19+ total B cells (C) in healthy 12-month-old FCG mice; at 10-week post BBN and at 1 week post 3$^{rd}$ BCG instillation in BBN exposed mice (BCG treatment initiated at week 7 post BBN initiation). Immune cell frequency was determined using multiparametric flow cytometry. Analysis of flow cytometry data was performed using FlowJo® software. GraphPad Prism was used to analyze the results and perform all statistical analysis. Statistical significance, *p<0.01, **p<0.001, ***p<0.0001, ****p<0.0001, was determined using two-way ANOVA with Tukey's post-hoc tests. Data represents mean ± SD. Representative multiplex IF stained bladder sections showing TLS and immune cell infiltration post 3rd BCG instillation in 12-month-old FCG mice (D; 200x magnification).

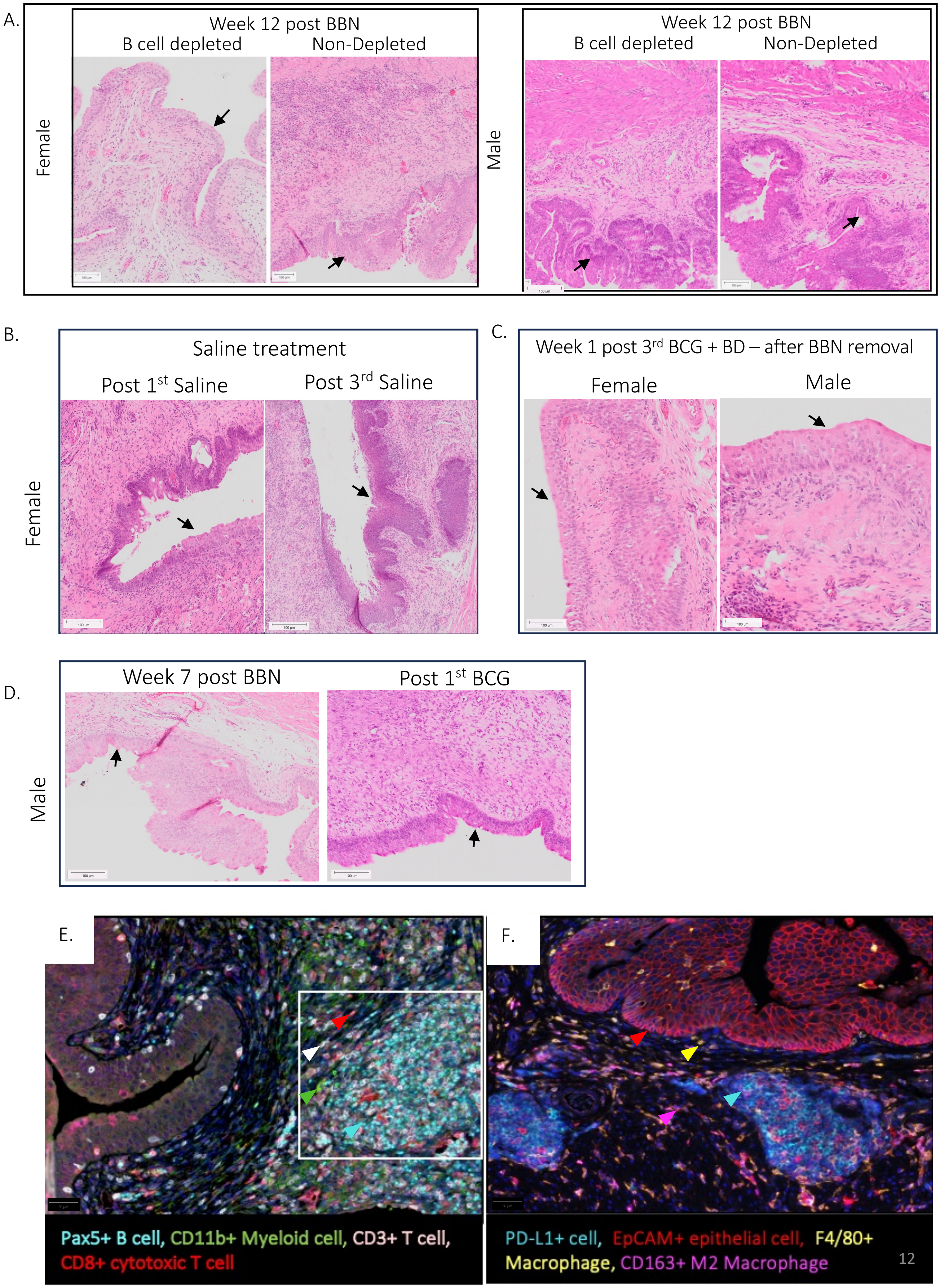
A.
Week 12 post BBN
B cell depleted
Non-Depleted
Female
Week 12 post BBN
B cell depleted
Non-Depleted
Male
B.
Saline treatment
Post 1st Saline
Post 3rd Saline
Female
C.
Week 1 post 3rd BCG + BD – after BBN removal
Female
Male
D.
Week 7 post BBN
Post 1st BCG
Male
E.
Pax5+ B cell, CD11b+ Myeloid cell, CD3+ T cell,
CD8+ cytotoxic T cell
F.
PD-L1+ cell, EpCAM+ epithelial cell, F4/80+
Macrophage, CD163+ M2 Macrophage

**Figure S3. Effect of transient B cell depletion and BBN exposure on urothelium of BBN exposed female and male mice.** Histopathological changes in the urothelium at 12 weeks post BBN exposure with or without B cell depletion in female and male mice are shown in the H&E-stained whole bladder section **(A).** In female mice, H&E-stained whole bladder section reveals histopathological changes in the urothelium at 1 week post 1$^{st}$ and 3$^{rd}$ Saline treatment with continuous BBN exposure **(B).** H&E-stained whole bladder section showing histopathological changes in the urothelium at 1 week post 3$^{rd}$ BCG + B cell depletion treatment after removal of BBN exposure in female and male mice **(C).** Histopathological changes in the urothelium at 7 weeks post BBN exposure and 1 week post 1$^{st}$ BCG treatment in male mice are shown in H&E-stained whole bladder sections **(D).** n=2-9 mice/group. Scale bar in H&E-stained images– 100 µm. Arrows indicate the urothelium. Representative multiplex IF stained bladder sections showing TLS and immune cell infiltration post 3rd BCG instillation (**E and F**; 200x magnification).

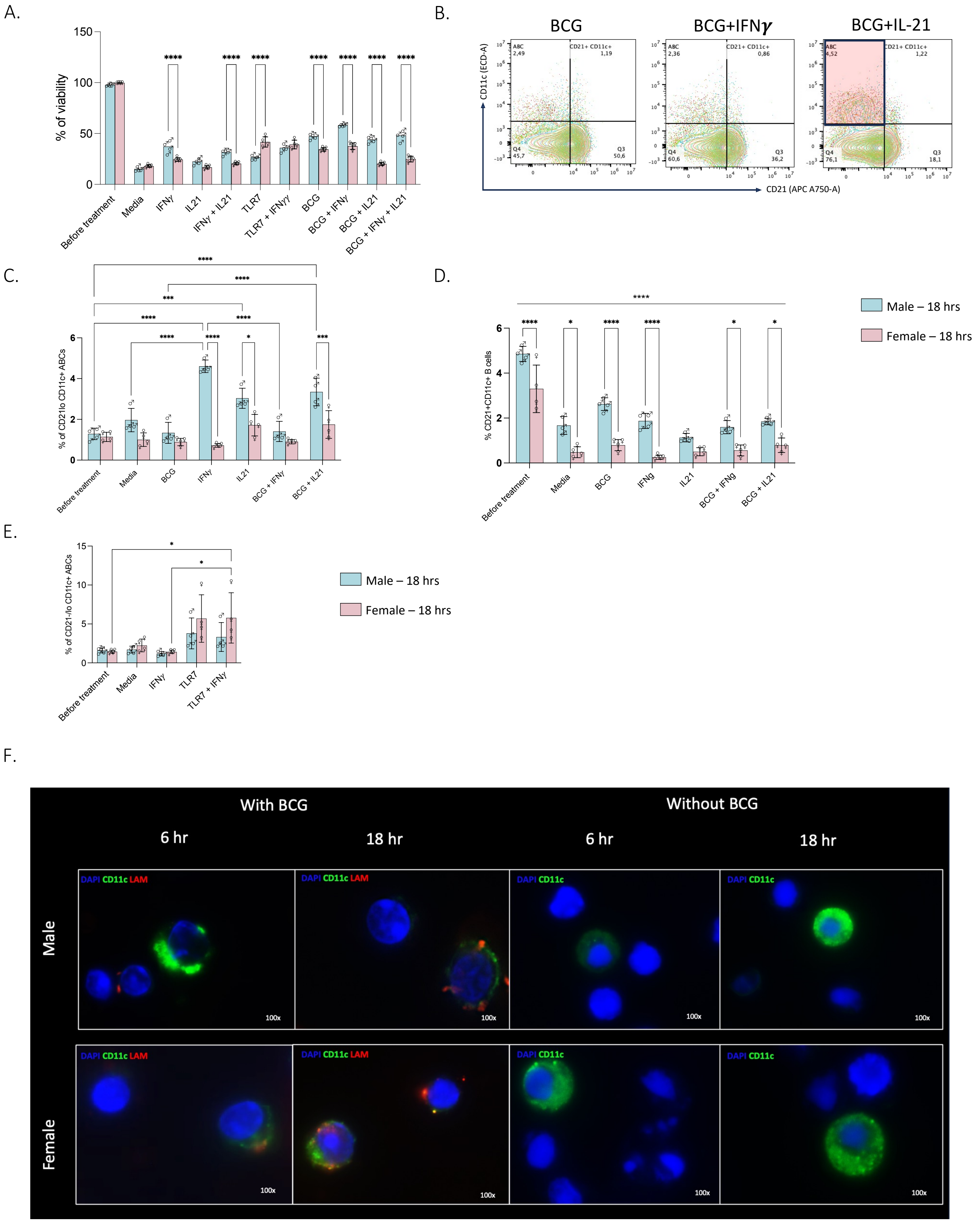
A.
% of viability
Before treatment
Media
IFNγ
IL21
IFNγ + IL21
TLR7
TLR7 + IFNγγ
BCG
BCG + IFNγ
BCG + IL21
BCG + IFNγ + IL21
B.
BCG
BCG+IFNγ
BCG+IL-21
CD11c (ECD-A)
CD21 (APC A750-A)
ABC 2,49
CD21+ CD11c+ 1,19
Q4 45,7
Q3 50,6
ABC 2,36
CD21+ CD11c+ 0,86
Q4 60,6
Q3 36,2
ABC 4,52
CD21+ CD11c+ 1,22
Q4 76,1
Q3 18,1
C.
% of CD21lo CD11c+ ABCs
Before treatment
Media
BCG
IFNγ
IL21
BCG + IFNγ
BCG + IL21
D.
% CD21+CD11c+ B cells
Before treatment
Media
BCG
IFNg
IL21
BCG + IFNg
BCG + IL21
Male – 18 hrs
Female – 18 hrs
E.
% of CD21-/lo CD11c+ ABCs
Before treatment
Media
IFNγ
TLR7
TLR7 + IFNγ
Male – 18 hrs
Female – 18 hrs
F.
With BCG
Without BCG
6 hr
18 hr
6 hr
18 hr
Male
Female
DAPI CD11c LAM
DAPI CD11c
100x

**Figure S4. B cell differentiation to ABCs following *in vitro* treatment with IFN-$\gamma$, IL-21 and BCG is sex-dependent.** Splenic B cells were isolated from 12-month-old male and female mice exposed to 7 weeks of BBN. Bar graph showing percentage of viability before and after 18 hrs of *in vitro* treatment with IFN-$\gamma$,IL-21, TLR7 and infection with BCG alone and in combination (as indicated) **(A)**. Contour plot showing spread of ABC population after IFN-$\gamma$ and IL-21 treatment and infection with BCG (increase in ABC population is shown with a red box in BCG + IL-21 treatment contour plot (**B**). Bar graph illustrating sex differences in B cell differentiation to ABC subsets B220+CD19+CD21lowCD11c+ **(C)** and B220+CD19+CD21+CD11c+ (**D)**, after 18 hrs of treatment with IFN-$\gamma$, IL-21 with or without BCG infection. Bar graph showing frequency of ABCs (B220+CD19+CD21-/lowCD11c+) **(E)** population after 18 hrs of treatment with IFN-$\gamma$ and TLR7. Analysis of flow cytometry data was performed using FlowJo® software. All statistical analysis was performed using GraphPad Prism. Ordinary two-way ANOVA with Tukey's post-hoc test was applied to determine statistical significance in differences (*p<0.05, **p < 0.01, ***p <0.001,****p<0.0001, —— indicates significance across all the treatment group with p<0.0001). Data represents mean ± SD. Immunocytochemistry of isolated B cells after 18hrs of treatment with IFN-$\gamma$, IL-21 and BCG using anti-LAM (**red**) and anti-CD11c (**green**) antibodies to identify *Mycobacterium bovis* and CD11c expression, respectively **(G;** 100x magnification)**.** DAPI (**blue)** was used to identify the cell nuclei.

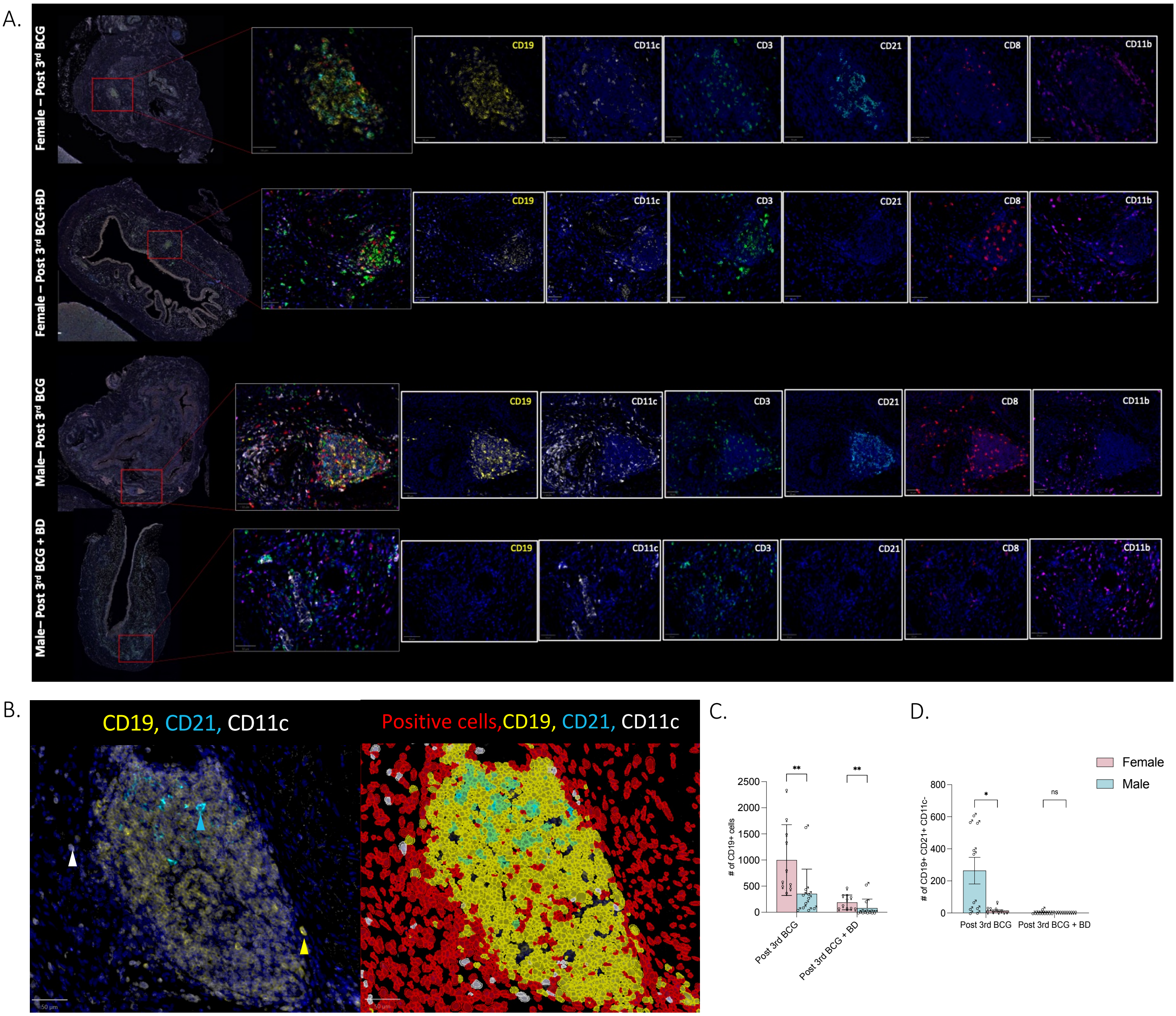


**Figure S5. Immune infiltration in the bladder microenvironment after repeated BCG treatment and B cell depletion in BBN exposed mice.** Multiplex IF stained bladder sections showing increased immune cell infiltration 1 week post 3rd BCG instillation with and without B cell depletion in 12-month-old female and male mice (**A**; 200x magnification). Representative multiplex IF stained bladder section showing positive cell detection and composite object classifier for identifying total B cells (CD19+), CD19+CD21-CD11c (ABCs), CD19+CD21+CD11c+ and CD19+CD21+CD11c- B cells **(B,** 200x magnification). Bar graph (mean ± SD) represents number of total CD19+ B cells **(C)** and CD19+ CD21+ CD11c- B cells **(D).** 10 annotations per treatment group used for the analysis. Cell detection analysis and composite object classifier was performed using QuPath software. All statistical analysis were performed using GraphPad Prism. Mann-Whitney test was applied to determine statistical significance in differences (*p<0.05, **p<0.001, ns – not significant).

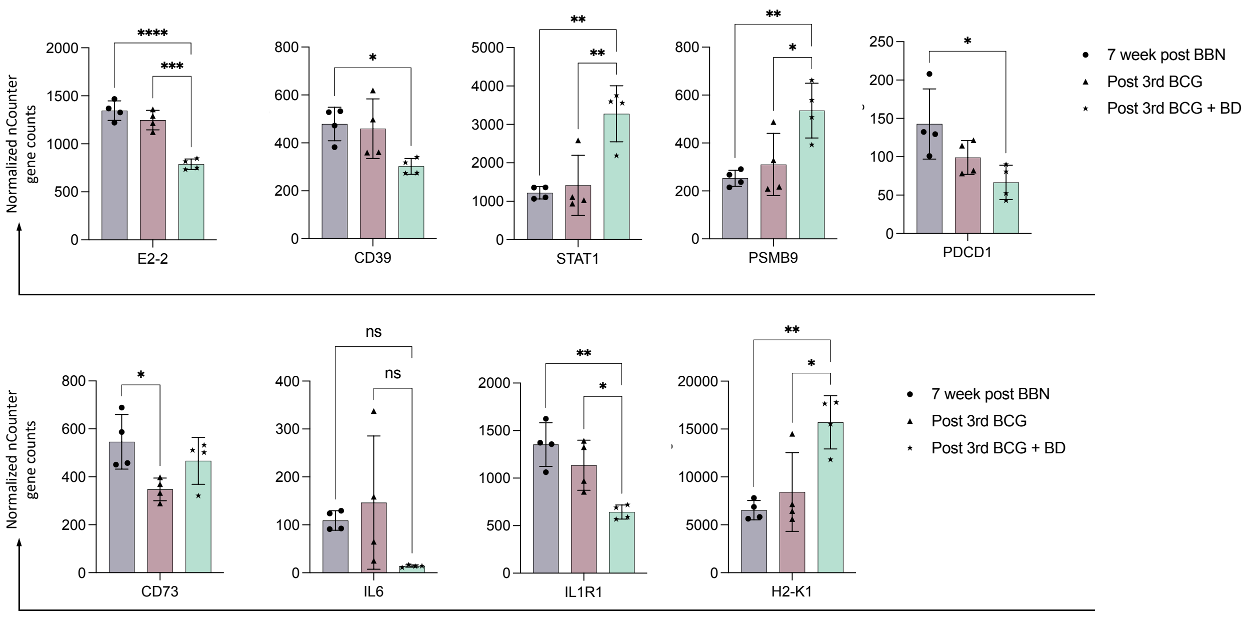


**Figure S6. B cell depletion alters expression profiles of immune regulatory genes in the bladder microenvironment.** Bar graphs illustrating the mean ± SD of normalized nCounter gene counts, analyzed using NanoString nSolver software, in three groups of female mice at 7-week post BBN exposure, one week post 3rd BCG with or without B cell depletion. Statistical analysis was conducted using GraphPad software with a one-way ANOVA test to identify significant differences in expression profiles across the various genotypes (*p<0.05, **p<0.01,***p<0.001, ****p<0.0001 ns – not significant). N=4 per group.

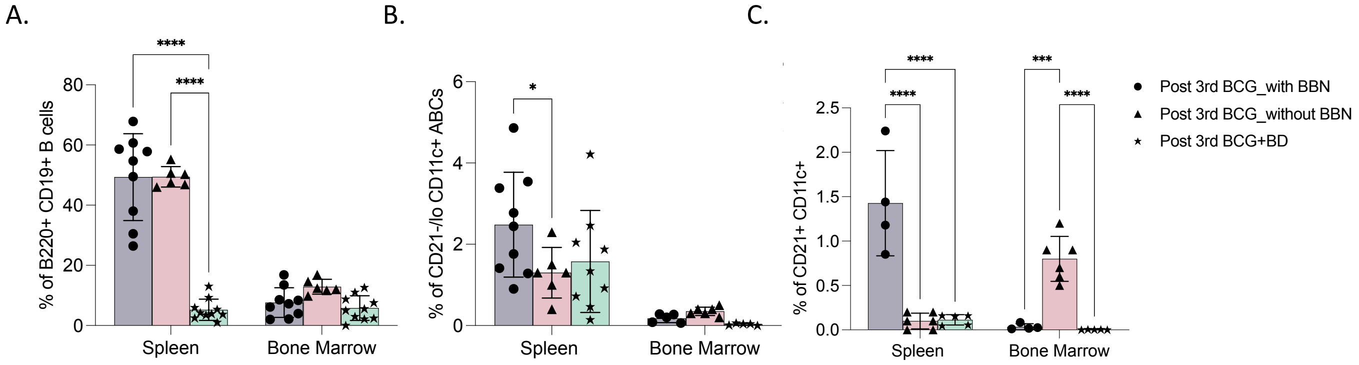


**Figure S7. Splenic ABC expansion is enhanced by a combination of BBN exposure and BCG treatment.** Bar graph (mean ± SD) represents the frequency of total B cells (B220+ CD19+ **(A)**), ABCs (CD21-/lo CD11c+ **(B)**) and CD21+ CD11c+ B cells **(C)** in 12-month-old female mice after 3 weekly instillations of BCG without BBN exposure to determine the role of BBN carcinogen in driving the expansion of ABCs. Analysis of flow cytometry data was performed using FlowJo® software. All statistical analysis was performed using GraphPad Prism. Ordinary two-way ANOVA with Tukey's post-hoc test was applied to determine statistical significance in differences (*p<0.05, **p < 0.01, ***p <0.001,****p<0.0001).

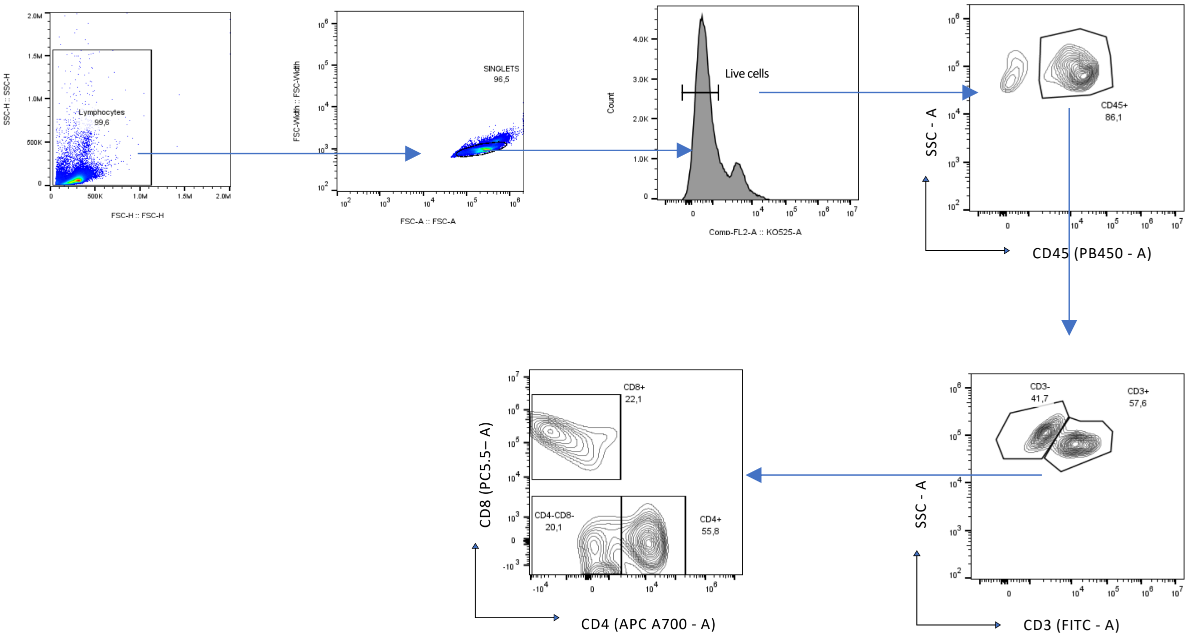


**Figure S8. Gating strategy for identifying different subsets of T cells.** Single cell suspensions from spleen and bone marrow of mice were stained with T cell panel antibodies and subjected to multispectral flow cytometric analysis. Single cells was first identified by plotting forward scatter width (FSC-Width) against side scatter (SSC-A). Events deviating from the linear cell population, called as doublets were excluded. Live cells were evaluated by gating on viability dye-negative cell. CD45+ marker was used to discriminate the total immune cell population and further used for identification of different subsets: CD3+ total T cells, CD3+ CD4+ CD8- T cells, CD3+ CD4- CD8+ T cells and CD3+ CD4- CD8- T cells. Single color (SC) and fluorescence minus one (FMO) was used to determine the position of gating for all subsets and further compared with unstained samples. All analysis were done in FlowJo® software.

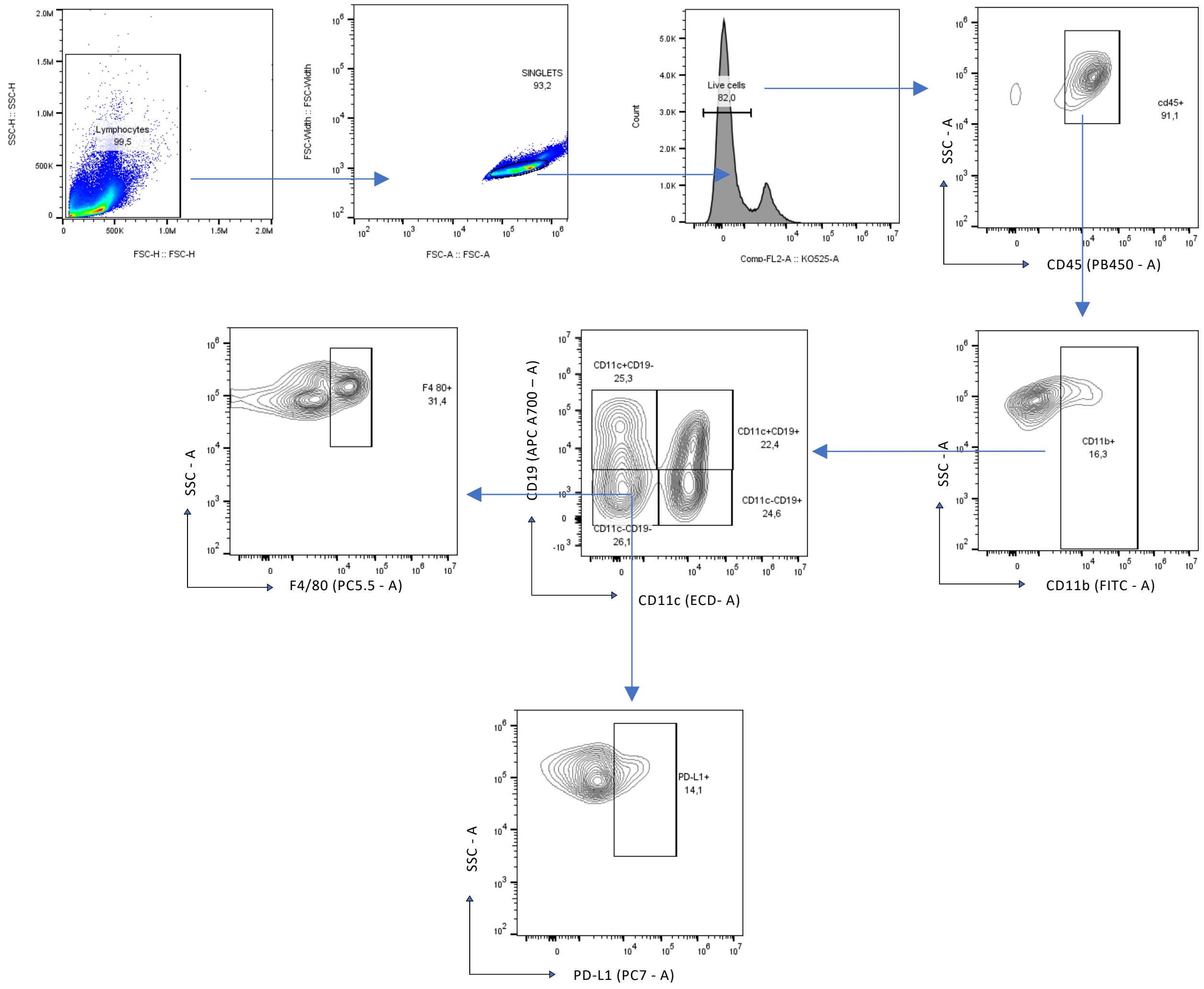


**Figure S9. Gating strategy for identifying myeloid cell population**. Single cell suspensions from spleen and bone marrow of mice were stained with antibodies and subjected to multispectral flow cytometric analysis. Single cells was first identified by plotting forward scatter width (FSC-Width) against side scatter (SSC-A). Events deviating from the linear cell population, called as doublets were excluded. Live cells were evaluated by gating on viability dye-negative cell. CD45 marker was used to discriminate the total immune cell population and further used for identification of different subsets: CD11b+ CD11c- CD19- myeloid cell, CD11b+ PD-L1+ cells and CD11b+ F4/80+ cells. Single color (SC) and fluorescence minus one (FMO) was used to determine the position of gating for all subsets and further compared with unstained samples. All analysis were done in FlowJo® software.

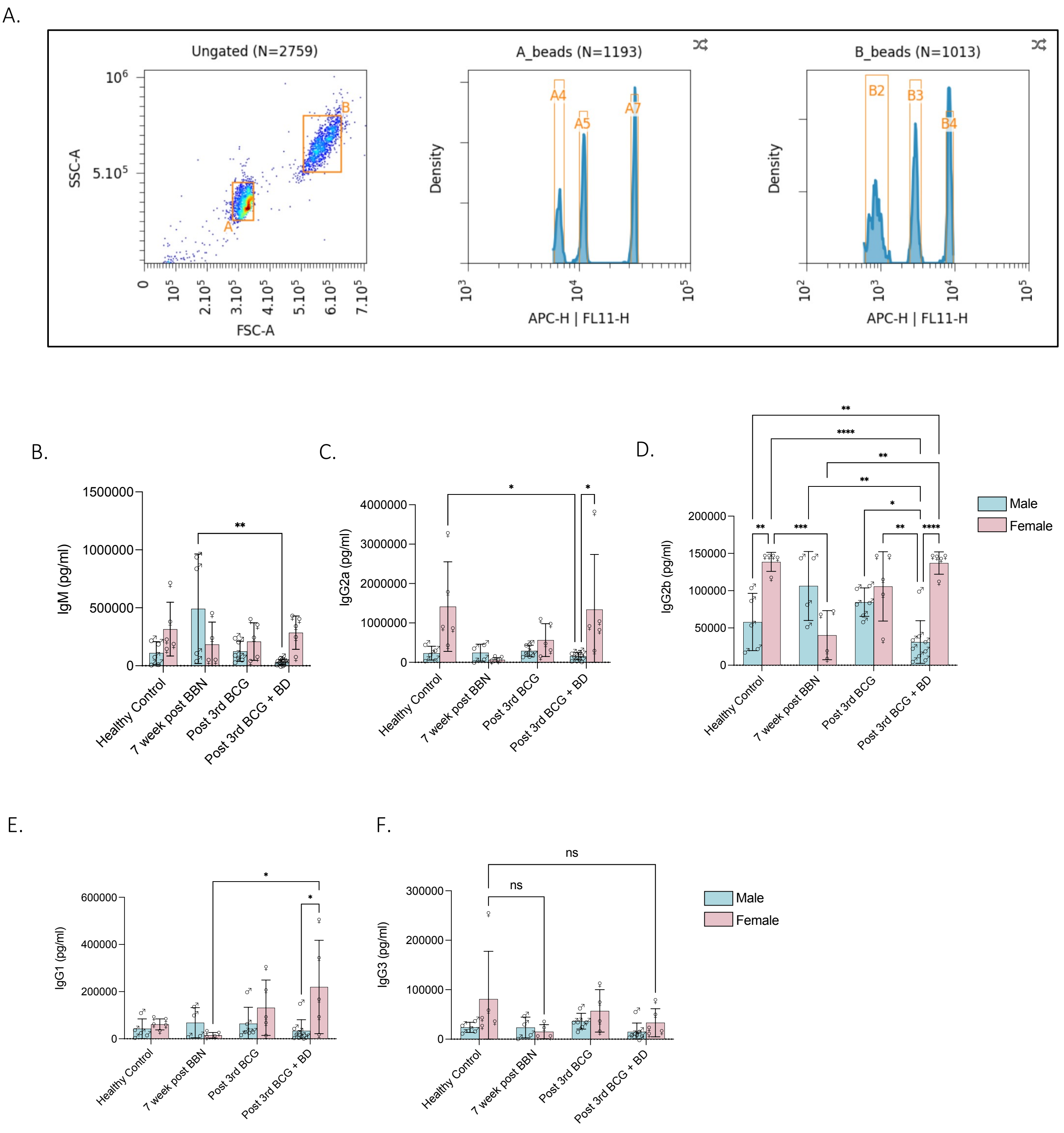


**Figure S10. BBN exposure and BCG treatment alters plasma immunoglobulin profiles in a sex differential manner.** Plasma collected at 1 week post 3rd BCG dose with or without B cell depletion was subjected to LEGENDplex Immunoglobulin isotyping assay and analyzed via flow cytometry. Data analysis were done using LEGENDplex Data Analysis Software Suite Qognit. Representative pseudo-colour plot and histogram illustrating clear separation between beads and gating of different immunoglobulin isotypes (6-plex) (**A**). Plasma from both males and females depicted differential profiles of plasma immunoglobulins indicative of B cell expansion and class switching (n=3-9 in each group, **B-F**). All statistical analysis was performed using GraphPad Prism. Statistical significance, *p<0.01, **p<0.001, ***p<0.0001, ****p<0.0001, was determined using two-way ANOVA with Tukey's post-hoc tests.
